# Spatial heterogeneity influences phage-host co-existence dynamics and phage resistance development in *Vibrio anguillarum*

**DOI:** 10.1101/2024.12.28.630609

**Authors:** Ling Chen, Mathias Middelboe, Sine Lo Svenningsen

## Abstract

Exposure of bacterial populations to phage infection pressure is a significant driver of phage-host co-evolution and diversification. However, the impact of the spatial organization of bacterial population ‘co-existence and co-evolution dynamics remains poorly understood. Here, we investigated how the spatial structure of the host population affects phage-host interactions by co-cultivating a *Vibrio anguillarum* strain with the lytic T4-like vibriophage KVP40 under conditions that resulted in either a homogenous, well-mixed population or a heterogeneous population with bacterial aggregates. We observed markedly different temporal dynamics associated with the two population structures over a 30-day adaptive laboratory evolution experiment. Phage and host dynamics suggested that phage-sensitive subpopulations retained in the aggregates substantially prolonged the coexistence of phages and bacteria, relative to the homogenous environment. However, the limited phage propagation on the sensitive subpopulations was insufficient for host range expansion to occur. In contrast, daily supplementation of sensitive host bacteria in parallel experiments readily led to the emergence of phage mutants with enhanced infective capability and an expanded host range. These results underscore the importance of bacterial spatial organization in determining the outcome of phage attacks and highlight the complex interplay between genomic processes and ecological conditions in driving evolutionary innovations.

## Introduction

Bacteriophages are the most numerous entities across Earth’s biosphere and control microbial communities through predatory, and sometimes mutualistic, interactions ^1–5^. A successful phage infection depends on molecular interactions with the bacterial host, in particular host binding though receptor recognition, why phages display a narrower host range than most other microbe predators ^6–8^. This is one reason for the renewed interest in utilizing phages as a more discriminatory alternative to antibiotics ^9–13^. Like antibiotics, however, phage infections impose a strong selection pressure for resistant subpopulations to arise and take over the bacterial population ^14–16^. Any phage mutants with the capacity to overcome host resistance then gain an advantage and can initiate another round of antagonistic interactions ^17–28^. This ongoing reciprocal adaptation shapes the dynamic interactions between phages and their bacterial hosts and fuels their genetic divergence, serving as a fundamental driving force for the evolution of wild microbial populations ^2,21^.

Reproducing phage-bacterial reciprocal adaptation cycles in adaptive laboratory evolution experiments has proven challenging. Phage-bacteria encounter rates are high and uniform in well-mixed monoculture experiments, resulting in a strong selection pressure for phage-resistant hosts. This stands in contrast to natural settings, where phage-bacterial encounters often occur within spatially heterogeneous environments and in highly diverse and sometimes dense communities with other microorganisms, resulting in complex evolutionary dynamics ^6,7,29–31^. Studies that include long-term monitoring of phage-bacteria coevolution in spatially heterogeneous environments are rare. Consequently, the effects of community composition and a spatially structured environment on focal phage-bacterial coevolution are not well understood ^30–32^, which limits our understanding of how phage predation shapes the diversity and drive the evolution of wild microbial populations.

In this study, we observed that simple alterations to the growth conditions of *Vibrio anguillarum* isolate PF4 ^33^ consistently resulted in two distinct population structures. When grown in glass flasks, *V. anguillarum* formed a well-mixed population of individual planktonic bacteria. In contrast, growth in glass tubes led to a spatially structured bacterial population, comprising a heterogenous mixture of large bacterial aggregates and individual planktonic cells. To explore how these bacterial physiological responses influenced the co-evolutionary dynamics between the bacteria and a lytic phage, we conducted long-term coevolution experiments. In summary, we monitored the evolution of the *V. anguillarum* isolate both in the absence of phages (control) and in the presence of an initially monomorphic population of the lytic vibriophage KVP40 ^34^ (coevolution). Our laboratory evolution experiments demonstrated that the spatially structured environment prolonged phage-bacterial coexistence substantially compared to the well-mixed environment, where phages were rapidly lost and phage-resistant bacteria prevailed. Our experiments also revealed that the prolonged phage-host coexistence in the spatially structured environment did not foster the evolution of phage host-range mutants, presumably due to low rates of phage replication, despite the presence of subpopulations of phage-susceptible cells in the bacterial aggregates. Consistent with the arms-race theory, however, we readily identified phage host-range mutants in cultures with abundant phage replication, facilitated by the daily addition of phage-susceptible cells.

To explore the impact of a multispecies context on phage-bacterial interactions, we included seven non-host strains that are phylogenetically divergent at the genus or even phylum levels, alongside the phage-host pair. In this complex setting, phages were rapidly lost, but phage-susceptible variants soon outcompeted the resistant mutants in the absence of phages. Thus, introducing a multispecies environment appeared to increase the fitness cost of phage resistance mutations in *V. anguillarum*.

## Results

### Minor variations in growth conditions lead to divergent population phenotypes

We first observed that when inoculated in rich medium, *Vibrio anguillarum* strain PF4 displayed divergent phenotypes in response to the specific growth conditions. Aggregation occurred when 5 mL bacterial cultures were grown in 35 mL glass tubes at 28°C and moderate shaking speed (160 RPM), henceforth referred to as tube growth. In contrast, the bacterial cultures remained uniformly dispersed during growth of 10 mL cultures in 100 mL flasks at 30°C and higher shaking speed (180 RPM), henceforth referred to as flask growth. Figure 1 shows phase contrast microscopy images of the bacteria in aliquots of cultures grown in tubes (Fig. 1a, without phage) and flasks (Fig. 1b, without phage). The fraction of bacteria that were found in the aggregated and free-living phases in 24-hour cultures grown under the tube and flask conditions were quantified. We did not detect any cell aggregates in the cultures in flasks, where the cell density reached OD_600_ = 4.11 ± 0.13 (Fig. 1c, “-phage”). By contrast, growth in tubes resulted in 43% cells in aggregates as well as a ∼40% lower total cell density than in the flasks (OD_600_ = 2.58 ± 0.13) (Fig 1d, “-phage”).

**Fig. 1.**
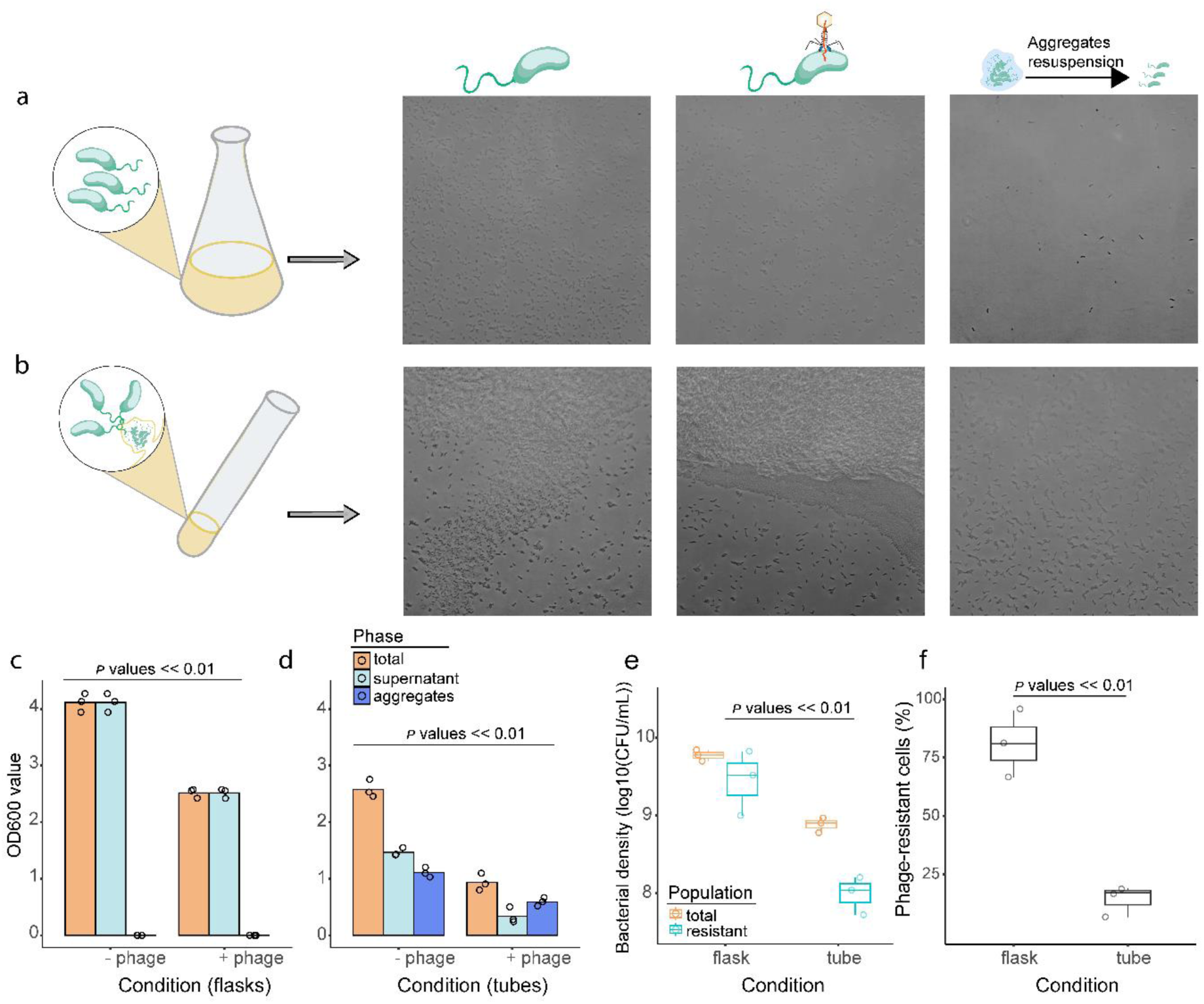
Different population structures of *V. anguillarum* PF4 cultures affect the accumulation of phage-resistant mutants. **a, b**, Morphological visualization of *V. anguillarum* PF4 aliquots after 24-hour culturing in flasks (**a**) and in tubes (**b**), in the absence (left) and presence (middle) of phage, and after resuspension of aggregates (right). **c, d**, Distribution of bacteria in the free-living phase and in aggregates when cultured in flasks (**c**) and tubes (**d**). Prior to enumeration, bacterial aggregates were resuspended as described in Methods. “+phage” refers to populations after 24-hours of growth in the presence of phage KVP40. **e**, Total bacterial concentration (CFU/mL) and the concentration of phage-resistant bacteria (CFU/mL) after overnight growth in the presence of phage KVP40, in flasks and tubes, respectively. **f,** Percentage of total population that is resistant to phage KVP40 after 24 hours growth in the presence of phage, in flasks and tubes, respectively. Values in panel **f** were calculated based on the data shown in panel **e**. *P*-values were determined by two-sided Wilcoxon tests with Dunn’s correction for multiple comparisons.

### Aggregation-prone tube growth reduced the accumulation of phage-resistant mutants

Having established the growth conditions for consistently obtaining homogeneous versus heterogeneous populations of *V. anguillarum* PF4, we investigated the effects of the two growth modes on phage infection. Cultures were co-inoculated with *V. anguillarum* PF4 and phage KVP40 and cultured for 24 hours in each of the two conditions described above. Addition of phage resulted in a lower bacterial yield under both growth conditions (Fig. 1c-d). While the phage had no effect on aggregation in the flasks (Fig. 1c, Supplementary Fig. S1a,c), addition of phage to the tube cultures increased the percentage of cells found in aggregates to 55%, (Fig. 1d, Supplementary Fig. S1b,d). In addition, while ∼80% of the cells that survived phage treatment in flasks were phage-resistant mutants, less than 25% of the surviving cells in tubes were resistant to phage infection (Fig. 1e-f). In line with a previous finding ^33^, this suggested that aggregation functions as a bacterial physiological defense mechanism providing a spatial sanctuary for the phage-sensitive *V. anguillarum* subpopulation.

### The coevolutionary dynamics between phage and bacteria are shaped by the spatial structure of the bacterial population

Due to the dramatic difference in accumulation of phage-resistant mutants after a 24-hour growth cycle in tubes versus flasks, we speculated that long-term phage-bacterial coexistence dynamics might also differ substantially between the well-mixed and the aggregation-prone growth conditions. To study coexistence dynamics of phages and hosts we propagated a set of replicate cultures under each growth condition for 30 days, by diluting the outgrown cultures 1:100 into fresh medium every 24 hours. The cultures were infected with phage KVP40 on Day 0. A set of control cultures without phage addition were processed in parallel. Phage and bacterial densities were quantified on select days and culture aliquots were stored for subsequent investigation. As shown in Figure 2, the two growth conditions resulted in two divergent coevolution trajectories. In terms of the overall bacterial population, the pattern was similar under both conditions, in that phage infection initially strongly suppressed bacterial growth, marked by a lower bacterial density in phage-treated cultures than in the control cultures on day 1 (Fig. 2a, d). A corresponding initial increase in the phage density was also observed under both conditions (Fig. 2b, e). Yet, in both environments, the bacterial population eventually recovered, and by the third time point it reached a density close to the control cultures without phage (Fig. 2a, d). At this point, most bacteria were resistant to the ancestral phage, as measured by randomly picking hundreds of colonies for each time point within each group and evaluating their susceptibility to the ancestral phage KVP40 by the cross-streaking method (Fig. 2 c, f).

**Fig. 2.**
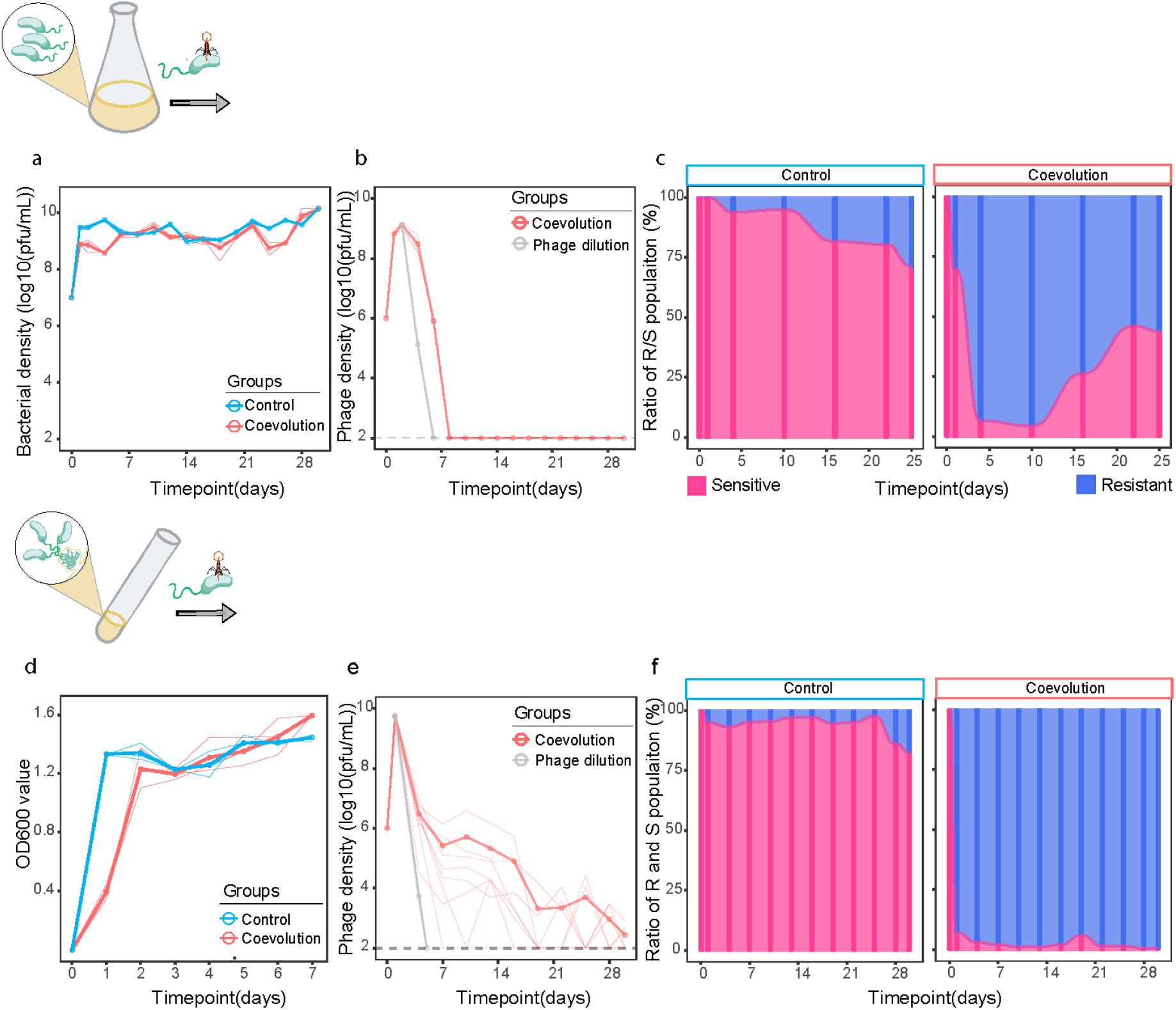
Dynamics of phage titer, bacterial population, and population composition under different conditions. **a**, **b**, Population dynamics of bacteria (**a**) and phage (**b**) during 30 days of serial culturing in flasks. The co-evolution experiments containing both phage KVP40 and *V. anguillarum* PF4 are shown in red (n = 3), while control groups without phage are in blue (n=3). **c**, proportion of bacterial colonies that were sensitive (pink) and resistant (blue) to the ancestral phage KVP40 at selected time points during growth in flasks (n = 120 isolates per time point). **d**, **e**, population dynamics of bacteria (**d**) and phages (**e**) during 30 days of co-culture in tubes. For OD_600_ measurements, three out of the replicates were tracked for seven days for both the control and coevolution. The phage population was quantified for all eight replicates throughout the experiment. In all cases, measurements were made at the end of a 24h growth period. **f**, proportion of bacterial colonies that were sensitive (pink) and resistant (blue) to the ancestral phage KVP40 at selected time points during growth in tubes (n = 200 isolates per time point). Gray lines in **b** and **e**, indicate the theoretical rate of loss of phages due to daily 1:100 dilution of the culture, in the absence of any phage replication. Thick lines with points indicate the average values in **a**, **b**, **d** and **e**, while thin lines show individual replicates.

Despite the similar dynamics of the host populations, the growth conditions had a strong effect on the persistence of the phage populations. When co-cultured in flasks, the phage became undetectable by day 7 (Fig. 2b). In contrast, during co-culturing with the spatially heterogenous populations in the tube cultures, phages persisted until the end of the experiment, albeit at low levels (Fig. 2e). As is typical for coevolutionary dynamics, substantial divergence was observed between replicates (Fig. 2e, thin lines). On average, the phage density continually declined towards the threshold of detection (1.0×10^2^ PFU/mL) in our experiments (Fig. 2e, thick red line). Importantly, continuous phage propagation must have taken place in at least seven of the eight replicate tubes, as the phages produced by the initial outbreak would have been lost from the culture by day 5 due to the daily 100-fold dilution of the culture, as indicated by the gray “phage dilution” line.

In an effort to determine which of the differing growth parameters in tubes versus flasks prompted the stark difference in phage population dynamics, we conducted a series of 10-day passage experiments, where we swapped the specific growth conditions (temperature, shaking speed, culture volume, initial phage-bacterial ratio) between the tube and flask vessels. Our observations revealed that the actual vessel (tube or flask) correlated with the spatial structure of the population, independent of the specific inoculation parameters mentioned above. Tube growth consistently resulted in the formation of bacterial aggregates, and flask growth resulted in a homogeneous, planktonically growing bacterial population. As shown in Supplementary Figure S2, the tube and flask vessels also correlated with prolonged phage persistence, independent of the swapping of other growth parameters. Specifically phages persisted at moderate concentrations (≥10^6^ PFU/mL) for the duration of the experiments in the aggregation-prone tube cultures. In contrast, phage concentration declined steadily and became undetectable by day 10 in the planktonically growing flask cultures, irrespective of the specific inoculation parameters. These results suggest that phages were more readily lost in planktonic cultures than in aggregates, highlighting a critical role for the spatial structure of the bacterial population in phage-bacterial population dynamics.

We also quantified how the evolutionary dynamics of the phage-resistance trait varied depending on the growth condition in the 30-day adaptive evolution experiments. For the homogeneous flask cultures, the proportion of phage-sensitive bacteria started to rise as soon as phages were lost from the population (day 7) and reached the same abundance as the resistant cells by day 20 (Fig. 2c). By contrast, the ratio of phage-sensitive to phage-resistant bacteria remained very low for the remainder of the experiment in the tube cultures with the spatially structured environment (Fig. 2f). These results suggest that a selective pressure for loss of the phage-resistance trait exists in the absence of phage, which is outweighed by the predation pressure exerted by the small but persistent phage population in the heterogeneous tube cultures (10^3^-10^4^ PFU/mL 24-hour after dilution, Fig. 2e).

Surprisingly, we observed up to 20% phage resistance in the control cultures, which had not been exposed to phages (Fig. 2c, left), challenging the hypothesis that resistance is counter-selected in the absence of phage predation. This prompted us to assess the phenotypes of resistant isolates further.

### Phenotypic and genomic implications of phage resistance in *V. anguillarum*

We then explored the fitness costs associated with phage resistance by measuring the maximum yield (Fig. 3a), growth rate (Fig. 3b), and biofilm formation (Fig. 3c) of phage-resistant isolates from tube cultures in the absence of phages. In general, resistant isolates from every time point grew at reduced rates and resulted in a lower yield after overnight growth than the ancestral phage-sensitive *V. anguillarum* PF4. The maximum yield was reduced by 14%-20% (*p* value <0.001) relative to the wild type, as measured by OD_600_ (Fig. 3a). The capacity for biofilm formation was also generally reduced in the resistant mutants, the average of isolates from each time point ranged from 6% to 25% (*p* value <0.001) lower than wild type (Fig. 3c).

**Fig. 3.**
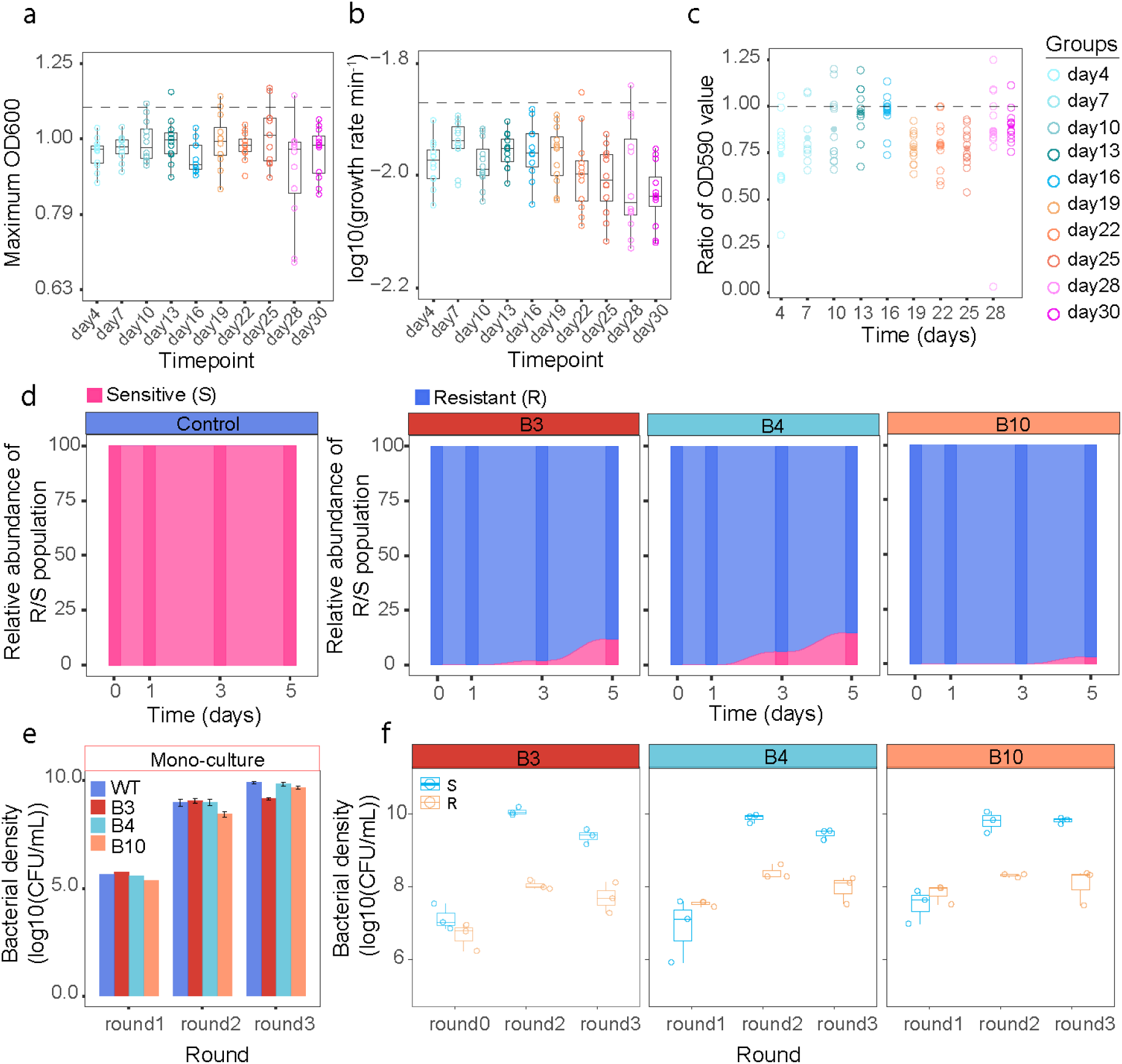
Characterization of diversified bacterial isolates from phage-bacterial coevolution in tubes. a, **b**, Maximum OD_600_ (**a**) and growth rate (**b**) of each resistant isolate relative to wild type PF4. The value of the wild type is indicated with a dashed line. Maximum OD_600_ and growth rate were determined from the OD_600_ values derived from growth curves. These curves were generated using three parameters (K: maximum OD_600_, N_0_: initial OD_600_ and r: intrinsic growth rate), which were fitted to the observed OD_600_ values using a logistic equation. **c**, The ability of each resistant isolate to form biofilm in the absence of phages. Each colored point indicates the mean value of each of the 12 isolates (with replicate n = 6). Values are normalized to the mean value of the wild type PF4. **d**, Proportion of sensitive and resistant bacteria in the populations on the indicated days, tested against the ancestral phage KVP40 (n = 30 isolates per time point). Three resistant variants, representing early (B3), middle (B4) and late (B10) time points of the coevolution experiment, were tested. The wild type host PF4 was included for reference. **e, f**, competition culture experiment individually (**e**) and paired with the wild type host PF4 (**f**). Experiments were performed for three rounds and each round was inoculated with the previous round’s end-point culture. The colony forming units (CFU/mL) of two phenotypes (phage-resistant and phage-sensitive) in the mixture were quantified as the subpopulation surviving a high titer of the ancestral phage, and the subpopulation killed by the ancestral phage (colonies formed in the absence of phage minus the colonies formed in the presence of KVP40), respectively. The *P*-values are determined by two-sided Wilcoxon test with Dunn’s correction for multiple comparisons.

To investigate if the fitness costs of phage resistance resulted in rapid mutation to phage sensitivity in the absence of phage, three resistant isolates obtained from different stages of the coevolution experiment were grown in the absence of phage in a 5-day transfer experiment. The result showed that while the phage-sensitive wild type strain maintained full susceptibility to the ancestral phage KVP40, and the majority of the resistant populations remained resistant for five days in the absence of phage, a small fraction of all three resistant populations reverted to phage sensitivity after 3-5 days (Fig. 3e). This confirmed that resistance to KVP40 is a reversible trait in *V. anguillarum* PF4. Next, we examined how the three phage-resistant variants performed in direct growth competition with the ancestral wild type strain in the absence of phages. A 1:1 mixture of cells from a resistant isolate and wild type *V. anguillarum* PF4 cells was cultured in rich medium to mid-exponential phase (round 1). The mixture was back-diluted into fresh medium and cultured to late exponential phase two additional times (round 2 and 3). The strains grew to comparable numbers in monoculture, except for the somewhat reduced growth of isolate B3 in round 3 (Fig. 3e). More importantly, the resistant isolates exhibited a markedly reduced density when co-cultured with the wild type strain, resulting in 10 to 60-fold lower densities after three rounds of growth competition (Fig. 3f). While this experiment did not allow us to distinguish wild type PF4 colonies from phage-sensitive revertants of the resistant isolates B3, B4 and B10, it demonstrated strongly reduced competitiveness of phage-resistant bacteria compared to the phage-sensitive counterparts. Overall, these results show that resistance to phage KVP40 comes with a fitness cost in the absence of phage, and support the notion that phage-bacterial antagonistic interactions facilitate phenotypic diversification of bacteria ^23^.

To investigate the resistant isolates at the genome level, we sequenced the genomes of ten representative isolates, including the three isolates used in the competition experiments (B3, B4, B10). The isolates were picked from three time points; day 4, 16 and 28 of the coevolution experiment in tubes. Comparative genomics revealed 11 mutated loci across the ten phage resistant isolates (Table S1, Fig. S3). The gene encoding the nucleoside-specific channel-forming protein Tsx (formerly OmpK), which serves at the receptor for phage KVP40 ^35^, was mutated in all 10 isolates. Isolates B6, B9 and B10 carry a (−1) frameshift mutation near the end of the *tsx* coding region, isolates B2 and B4 carry a nonsense mutation two-thirds into the *tsx* coding region, isolate B7 harbors a 26-bp intragenic duplication in the middle of the *tsx* gene and isolates B1, B3, B5 and B8 carry a 90 bp deletion in the beginning of the *tsx* gene. These mutations are all consistent with the loss of protein function, therefore the *tsx* mutation is likely responsible for the phage-resistance phenotype in all 10 isolates.

Besides the phage receptor, three mutations were present in the majority of isolates: 9 out of 10 isolates (-B4) had a 1-2 bp insertion in *queG*, predicted to encode epoxyqueuosine reductase, an enzyme that catalyzes the tRNA modification queousine. Nine out of 10 isolates (-B5) shared two mutations, predicted to result in a C580S substitution in carbamoyl-phosphate synthase large chain protein, and a T307R substitution in glytathionyl-hydroquinone reductase (Table S1). The three gene functions are not intuitively related to each other, nor to the predicted nucleoside transport activity of *tsx*, and it is therefore not readily apparent whether they would affect the fitness of the phage-resistant isolates.

### Increased infectivity and lack of host range expansion in coevolved phages

To probe if the prolonged coexistence of phages and bacteria in tube cultures involved a classic arms race, where phage mutants with expanded host range were able to infect otherwise resistant bacterial isolates, we challenged 120 resistant bacterial isolates (12 per timepoint) from tube cultures with a total of 196 phage variants collected from the same tube cultures, to establish an all-by-all cross-infection matrix. The phages were isolated from culture aliquots using wild type *V. anguillarum* PF4 as the host. The results showed that none of the phage variants could form plaques on any of the bacterial isolates, besides the ancestral wild type (Supplementary Fig. S3). These results suggest that the prolonged presence of phages in the tube cultures is not due to host range expansion. Rather, phage propagation is likely maintained at low levels by the continued presence of phage-sensitive bacteria in the aggregates.

To investigate the effect of long-term phage-bacterial coexistence on phage genotypes, we sequenced the genomes of 10 phage isolates selected from the collection of 196 phages isolated from the 30-day coevolution experiment in tubes ((3 from day 4, 3 from day 16 and 4 from day 28). Conducting *de novo* genomic comparisons, we identified a total of 69 mutation sites across the 10 phages, most of which were located in genes with unknown functions (Table S2, Supplementary Fig. S4). There were 27 mutations in genes involved in the core replication machinery and phage structural genes. Of the structural genes, 6 were located on tail fiber subunits and all others were associated with head and neck passage structures (Table S2). Tail fiber mutations are frequently associated with host range expansion as they are involved in phage attachment ^36^, although in this case we did not identify any phages that could infect resistant variants of PF4 (Supplementary Fig. S3). Nevertheless, the detected mutations in the 10 phage isolates suggested an ongoing diversification and genomic divergence of phages during co-culture with the host (Supplementary Fig. S5).

We therefore compared key phenotypic traits, namely the adsorption and the maximum phage titer reached by the ten sequenced coevolved phages to that of the ancestral phage, using the wild type strain and the three representative resistant isolates, B3, B4 and B10 as hosts. As expected, the ancestral KVP40 phage adsorbed to the wild type host but not to the resistant isolates (Fig. 4a), confirming that the mutations in the *tsx* receptor gene in the resistant isolates had abrogated phage binding. The coevolved phage variants also showed no significant adsorption (*p* value > 0.05) to the resistant strains, confirming the lack of host range mutants demonstrated by the all-by-all infection matrix. However, one of the coevolved phages (Coevp10), showed 2-fold greater adsorption to the wild type host within 15 minutes than the ancestral phage (Fig. 4a), demonstrating an improved infectivity of the coevolved phage. This was further supported by 12-h infection experiments using the wild type strain as the host, in which two (Coevp9 and Coevp10) of the coevolved phages isolated from the later part of the 30-day coevolution experiment displayed a 60-fold higher phage titer than the ancestral phage (Fig. 4b, Fig. S6). To explore this phenotype, we examined infection by phage Coevp10 more closely (Fig. 4c). Coevolved phage Coevp10 indeed adsorbed more readily to the PF4 strain and had already increased its titer by 20 minutes after phage addition, while the ancestral phage remained in the adsorption phase throughout the 30 min experiment (Fig. 4c). These results demonstrate an overall faster infection cycle in the coevolved phage, which likely explains the higher phage titer achieved by this phage after 12 hours (Fig 4b, Fig. S6). In summary, these results demonstrated that the low-rate replication of a small phage population in the heterogeneous populations of bacteria grown in tubes supported the appearance of phage mutants with enhanced infectivity of susceptible host bacteria, while host range expansion mutants were not detected.

**Fig. 4.**
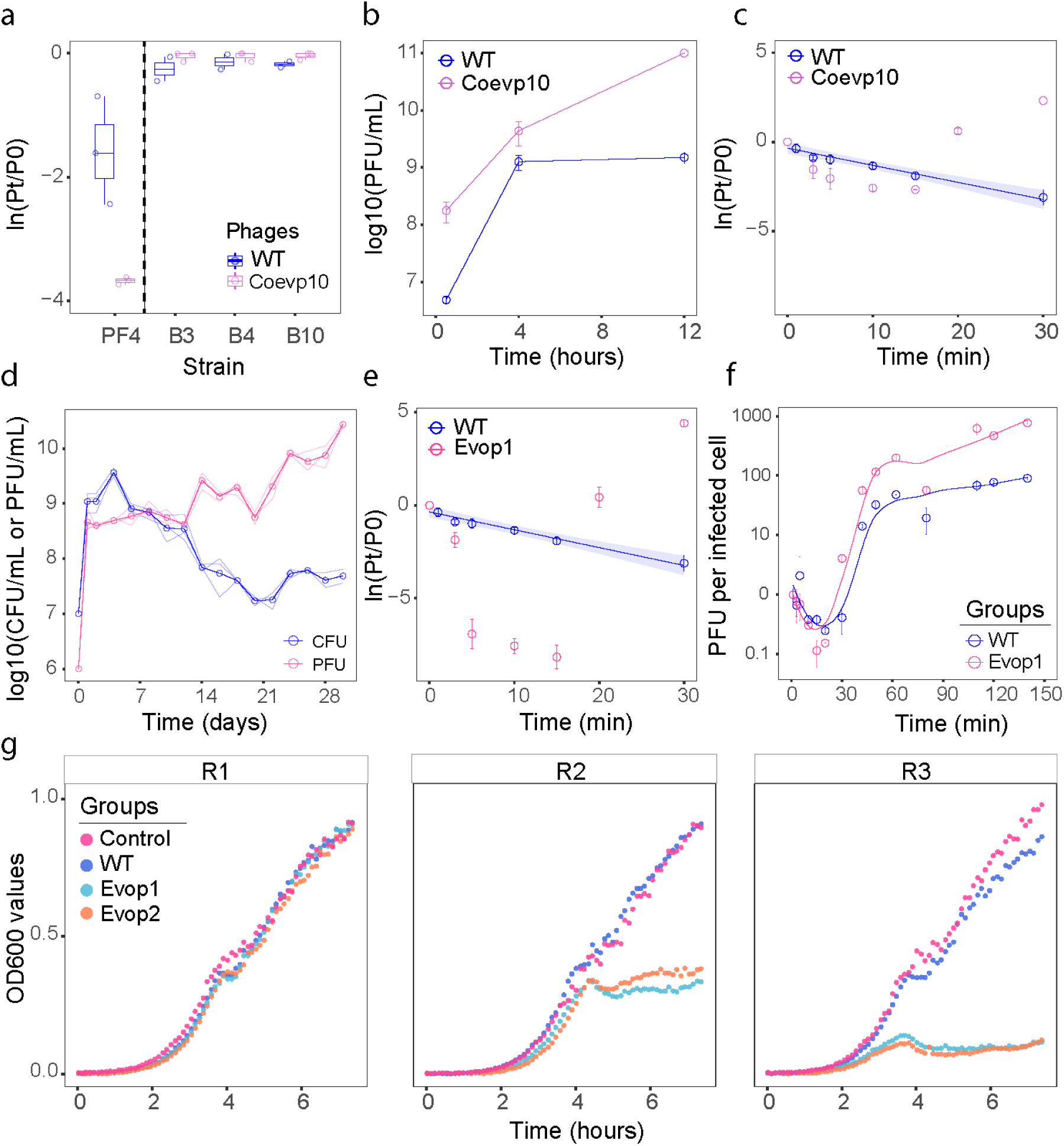
Infective traits of phage isolates from laboratory evolution experiments. **a**, Phage adsorption to PF4 and three resistant isolates (B3, B4, and B10) was quantified as the unabsorbed phages (PFU/mL) after an adsorption period (Pt) relative to the phage titer added at time 0 (P0). Pt = 30 min for the resistant bacterial strains. Pt = 15 min for strain PF4. **b**, **c**, Infective traits of the ancestral KVP40 and coevolved phage variant Coevp10, determined using the wild type PF4 host at times 0.5h, 4h and 12h after phage addition. **d**, Population dynamics of phages (PFU/mL) and surviving bacteria (CFU/mL) for a 30-day phage evolution experiment. Measurements of phage and bacterial concentrations were always made at the endpoint of a 24h cycle, immediately prior to dilution of phage in fresh bacterial culture. **e**, Adsorption curves of an evolved phage (Evop1) as well as wild type phage KVP40. **f**, One-step growth curve of Evop1 and wild type phage KVP40. **g**, Growth of three resistant bacterial isolates (R1, R2, R3) after addition of ancestral phage KVP40 (WT) and two evolved phage isolates (EvoP1, EvoP2). In the negative control, an equal volume of LM was added instead of phage lysate. The *P*-values are determined by two-sided Wilcoxon test with Dunn’s correction for multiple comparisons.

### Trained phages accumulated multiple mutations and acquired an increased bacterial predation efficiency

To enable comparison of our results to conditions supporting high rates of phage replication, we established phage-host co-culture experiments in flasks, where cell-free supernatants containing phage underwent daily transfer into cultures containing fresh susceptible hosts at every 24hr transfer, thus repeatedly exposing the phage population to wild type host bacteria. Figure 4d shows the dynamic changes in phage and bacterial concentrations over the course of 30 days of this phage evolution experiment. Phage and bacterial concentrations were always quantified at the end of a 24hr cycle, prior to inoculation of the phage in a culture of fresh bacteria. As the number of transfers increased, phages exhibited stronger predation of the bacterial population, resulting in lower densities of surviving cells in the cultures. In keeping with this observation, the phage population underwent a substantial increase in reproductive capacity, as the cultures yielded notably higher phage titers from transfer to transfer (Fig. 4d).

To begin to assess the effects of continued phage transfer on phenotypic divergence, we characterized phages from three plaques obtained on day 28 (Evop1-3). Compared to the ancestral KVP40 phage, these three phage isolates, which we refer to as trained phages, displayed a smaller plaque size on the wild type *V. anguillarum* PF4 host. Initial phenotypic characterization of the phage isolates showed that they displayed a much higher adsorption rate than the ancestral (WT) phage. More than 99.9% of the trained phages had adsorbed to host cells in the first five minutes, while the wild type phages had not adsorbed to the same extent even after 30 min (Fig. 4e, Supplementary Fig. S7). Furthermore, one-step growth of phage Evop1 demonstrated a 1.5-fold increase in burst size relative to the WT phage, significantly increasing the phage production rate of Evop1 (Fig. 4f). At the genome level, compared with phages from the coevolution experiment, the trained phages had a higher frequency of mutation sites. Most notably, the three phages shared a large, 5,711 bp, deletion (Table S2). Notably, the trained phage isolates showed an expanded host range. When they were mixed with three PF4 variants that were resistant to the ancestral KVP40 phage, they were able to infect two of them (Fig. 4g). Together, this demonstrates that trained phages outperformed the ancestral phage on multiple infection parameters, namely adsorption rate, burst size and host range.

### Increased cost of phage resistance in a mixed bacterial community

Finally, we assessed the effect of a mixed bacterial community on the specific phage-bacterial dynamics. We expanded our model system with seven non-host strains (Supplementary Fig. S8) forming a diverse microbial community. The mixed community was cultured in tubes for an 8-day long transfer experiment and the population dynamics of *V. anguillarum* PF4 in the presence of phage KVP40 was compared with parallel control cultures without phage.

Daily measurements of population densities revealed only minor differences in total bacterial density (Fig. 5a). Initially, *V. anguillarum* PF4 in the mixed community cultured with phage was clearly reduced relative to the cultures without phage (Fig 5c, day 1). However, phage-resistant *V. anguillarum* mutants rose to dominance after two transfers (Fig 5d, day 2), the phage population rapidly declined (Fig. 5b), and *V. anguillarum* PF4 were strongly represented again in the mixed cultures both with and without phages by day 4 (Fig. 5c). Interestingly, phage-sensitive *V. anguillarum* variants returned to dominate the *V. anguillarum* population in the mixed culture concurrent with the loss of the phage population (Fig. 5c and 5d, day 4). The rapid switch from resistant to sensitive bacteria after phages were lost from the cultures suggests that the fitness cost of resistance in the absence of phages is larger when *V. anguillarum* is growing in competition with other species (Fig. 5d) than when *V. anguillarum* is grown in monoculture (Fig 2c).

**Fig. 5.**
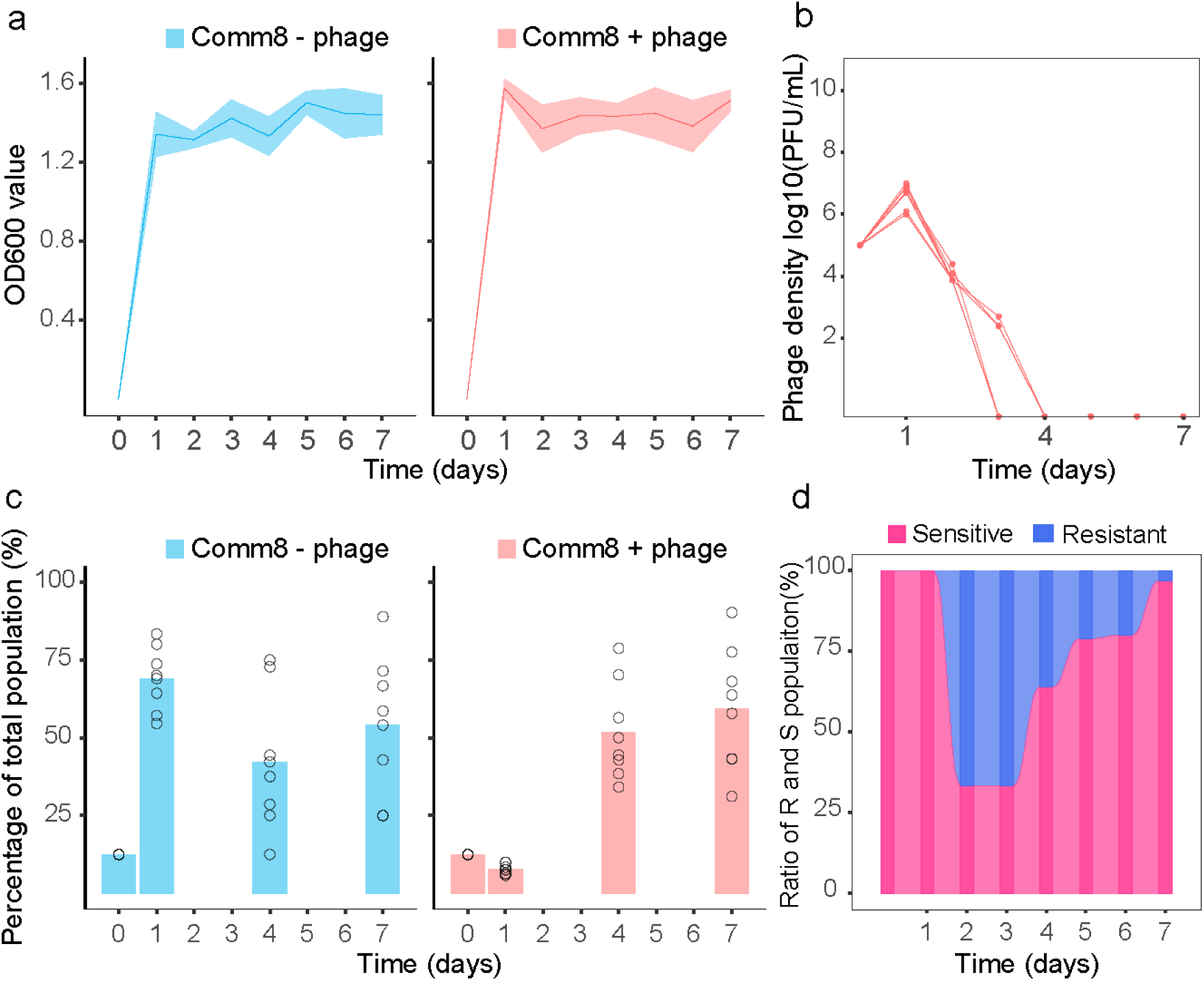
Dynamics of phage titer, bacterial population, and population composition in constructed multispecies consortia. **a**, Total absorbance of cultures of PF4 mixed with a community of seven other species (Comm8), in the presence or absence of phage KVP40. **b**, phage titer (PFU/mL) in supernatants harvested from Comm8+phage cultures. **c**, Proportion of colony-forming bacteria (CFU/mL) that are Vibrios at selected time points **d**, Proportion of Vibrios at selected days that were sensitive (pink) or resistant (blue) to the ancestral phage KVP40 (n = 30 isolates per timepoint). All measurements were made at the end of the indicated 24hr incubation cycle, before dilution of the culture into fresh medium.

## Discussion

In this study, we demonstrated that the co-existence dynamics of phage and bacteria is strongly influenced by the spatial organization of the bacterial population. Under laboratory conditions, we developed a model infection system comprised of a *V. anguillarum* strain and the lytic phage KVP40, in which a shift from homogenous (free-living cells) to heterogenous (free-living and aggregated cells in the same culture) population characteristics could be easily mediated by changing the culture vessel. By comparing tube and flask growth during 30-day phage-host adaptive laboratory evolution experiments, we revealed two divergent outcomes for phage-bacteria dynamics. In well-mixed flask populations, a rapid selection for phage-resistant bacteria and the loss of phages was observed. By contrast, phage-host coexistence throughout the experiment was observed when culturing heterogeneous populations. Similar observations of rapid phage extinction in well-mixed cultures were found for coevolution studies of Pseudomonas *fluorescens* SBW25 and lytic phage SBW25 Ф2, and *E. coli* with phage λ) ^37,38^, suggesting that in well-mixed lab monocultures there is such a strong selection for phage-resistant mutants that they quickly rise to dominance, cause phage proliferation to stagnate, thus reducing the probability of adaptive coevolution. In contrast, when co-cultured in environments with a heterogeneous spatial structure of the bacterial population, we suggest that the aggregates provided a refuge against phage infection which allowed a phage-sensitive host population to persist. This supports previous work with a related isolate of *V. anguillarum*, where cell aggregation was shown to represent a barrier for phage KVP40 infection ^33^. We observed that the continued presence of a sensitive host population enabled sufficient phage propagation to maintain a small phage population during long-term laboratory evolution experiments, in the current study lasting at least 30 days with daily dilution into fresh growth medium, which amounts to more than 200 bacterial generations. Our results provide evidence for the notion that spatial structure can alter phage-bacteria ecological interactions and impact the outcomes of phage-bacterial coevolution ^39^.

Several mutations were found in the genomes of phage-resistant variants. These included genes related to cell division, enzymatic functions, and outer membrane proteins. All sequenced isolates had mutations in the receptor gene (*tsx*/*ompK*) for the ancestral phage KVP40 ^35,40^. Such receptor mutations have been consistently identified as the key driver of resistance in laboratory coevolution studies ^8,23,25,27,41^. The role of the Tsx receptor in *V. anguillarum* physiology is not well understood, beyond its annotation as a nucleoside-specific, channel-forming, outer-membrane protein. It is therefore difficult to speculate if any of the accompanying mutations played a role in enhancing the fitness of the *tsx*^-^ phage-resistant mutants. The three tested phage-resistant isolates all performed poorly relative to the wild type in direct competition assays, illustrating that resistance to phage KVP40 is associated with fitness costs in *Vibrio anguillarum*, in agreement with general phage-host evolution theory ^21,42^. Such fitness costs have previously been reflected in impaired physiological properties ^28,43,44^, such as reduced growth rate and reduced biofilm formation as also identified in resistant clones obtained in this study. Consistent with previous reports ^38,45,46^, we observed that the phage-sensitive population increased following the loss of phages from the cultures. The fitness cost of resistance appeared to be amplified when competing bacterial strains were present in a multispecies setting, leading to an accelerated take-over of the population by phage-sensitive variants. The apparent high selection pressure for a functional Tsx receptor in more complex, multispecies environments could contribute to the selection for alternative non-mutational defense mechanisms, including downregulation of Tsx expression, putative abortive infection systems, biofilm formation and secretion of extracellular proteases that have been reported previously for *V. anguillarum* ^40,47^.

Even cultures started from a clonal population of phage-resistant variants readily accumulated phage-sensitive variants during short-term transfer assays in the absence of phages. Thus, reversible receptor mutations represent a somewhat dynamic and flexible response to phage exposure, which is in line with investigations of another fish pathogen, *Flavobacterium psychrophilum*, where reversible mutations in gliding motility and virulence genes were found to be flexible and efficient phage defense mechanism^48^. Despite the mutations to phage resistance frequently implying fitness costs ^25,42,49–51^, our study demonstrates how environmental parameters such as the presence of competing bacterial species, or the spatial structure of a monospecies population, can affect the balance of cost and benefit for resistance traits, thereby playing a key role in determining the coevolutionary dynamics of phage-bacteria as well as impacting the genetic divergence of bacterial populations ^43,44,52–56^.

Indeed, the impact of phage infection on the origin and maintenance of sympatric host diversity is largely dependent on host physiology and spatial heterogeneity ^57–61^. Since phages are obligate parasites, their distribution depends heavily on the distribution of bacterial hosts, regardless of the nature of the phage-host relationship, which spans a continuum from parasitism to mutualism ^62–64^. Therefore, by constraining the mobility of the hosts, e.g. by aggregation, the probability of encounters between phage and susceptible hosts are reduced, which impacts the outcome of phage predation ^59,60,65^. In addition, spatial heterogeneity allows for the formation of subpopulations and local adaptation, lowering the selective pressure by phages and promoting the coexistence of the bacterial host and their parasites^23,26,46,59,66–69^.

Genomic sequence data revealed that all phage isolates had divergent genomic variation, albeit with a tendency for higher similarity between phages harvested on the same day than between phages harvested on different days (Fig. S4). We note that phages Coevp9 and Coevp10, which both yielded increased phage titers, are almost genetically identical, varying only in one locus. Since most of the shared mutations are in hypothetical genes, it was not possible to speculate further on the genetic basis of the increased titers. The increased titer and adsorption of the coevolved phages is consistent with previous reports that long-term coexistence with bacteria confers phages with advantageous traits that significantly boosts their infectivity of the original host ^38,70^.

Interestingly, the trained isolates from the evolution assay where phage lysates were exposed to fresh susceptible hosts daily all shared a large fragment deletion containing mostly genes of unknown function. For myoviruses with large genome sizes, selective elimination of unnecessary genes under stress have been argued to benefit the progeny phages in future infections ^17,71–73^. We hypothesize that the reduced genome size of the evolved phages in this study may similarly contribute to the increased killing efficiency of the trained phages.

In general, long-term phage-bacterial coexistence requires adaptations and counter-adaptations in the sense that mutated phages with the ability to infect evolved hosts would be expected to arise. This is the fundamental theory of arm-race coevolution ^17,74^. Nevertheless, we did not identify host-range mutants in the coevolution experiments. By contrast, we readily isolated phage mutants with expanded host range among the phages that were trained daily on fresh susceptible hosts. In all likelihood, the important difference between the two scenarios is that many more rounds of phage replication take place between transfers in the training case than in the coevolution experiment, where the limited availability of susceptible hosts keep phage replication at a minimum. This observation supports that susceptible strains can be used as a stepping-stone to accelerate phage diversification, which in turn enhances the likelihood of infecting new hosts ^8,27,70,75^.

In summary, the divergent phenotypes of *Vibrio anguillarum* PF4 cultures grown in tubes versus flasks comprised a convenient arena for studying the effects of bacterial aggregation on phage-bacterial coevolution. Taken together, our results demonstrate that spatial structure of the bacterial population and the variable, environment-specific fitness costs of phage resistance traits play pivotal roles for phage-host population dynamics and hence phage-bacterial coevolution trajectories.

## Methods

### Bacteria and bacteriophage strains

*V. anguillarum* PF4 (accession id: CP010080/CP010081) and phage KVP40 were obtained in a previous study ^47,76^. Seven non-Vibrio strains were kindly provided by Zhipeng Huang (SIAT (www.siat.ac.cn)). Genomic sequences of all bacteria as well as the ancestral phage used in this work are available at NCBI. Experiments were conducted with bacteriophage KVP40 and *V. anguillarum* PF4, and in some cases together with seven other bacterial strains. Before the experiment, a single colony of each bacterial strain was cultured in LM (10 g/L tryptone, 5 g/L yest extract, 10 g/L sodium chloride in distilled water, autoclaved) at 30℃ with shaking unless otherwise noted. A KVP40 lysate was prepared by inoculating 100 µL of overnight growing *V. anguillarum* PF4 culture in 5 mL LM for 30 min and infecting with 100 µL of the original KVP40 stock until culture clearance was reached (2∼3 hours at 30℃). After centrifugation, the lysate was sterilized through a 0.22 um Millex-GV filter, supplemented with 0.1% chloroform, and stored at 4℃. Phage titer was estimated by quantitative plaque assay as described ^77^.

### Growth conditions in tubes and flasks

Cultures of *V. anguillarum* isolate PF4 were grown in either of two conditions, referred to as tube and flask growth, respectively. Tube growth refers to inoculation of an overnight culture into 5 mL of LM medium in 35 mL glass tubes and incubating at 28°C and moderate shaking speed at 160 rounds per minute (RPM) for 24 hours before sampling. Flask growth refers to inoculation of an overnight culture in 10 mL LM medium in 100 mL Erlenmeyer flasks and culturing at 30°C and high shaking speed (180 RPM) for 24 hours before sampling. Prior to phage infections, cultures were prepared as above and grown for 30 min, before two sets of cultures were infected by 10^7^ phage KVP40 shaking for 24 hours. At the end of the experiment, 1 mL of bacterial and phage mixture was collected for colony counts. To separate aggregates from planktonic cells, cultures were centrifuged briefly (200 RPM, 5s). The supernatant was separated from the pelleted aggregates, and aggregates were resuspended in the original volume of fresh medium using strong vortex; 5000 RPM, 10s. The biomass of each fraction was estimated by measuring OD_600_.

To measure the fraction of resistant cells, aliquots of 100 µL of a ten-fold dilution series of each replicate were spread on LM agar plates, as well as LM agar plates coated with 500 µL of 10^9^ PFU/mL KVP40 phage lysate. The fraction of resistant cells in the culture was estimated as the concentration of colony forming units (CFUs) on phage-coated plates (resistant population) relative to the concentration on no-phage plates (total population).

### Population composition in heterogeneous response to phage infection

To determine the effects of phage infection on population composition in tubes, the subpopulations in supernatants and aggregates were assessed individually. Overnight cultures of PF4 were diluted 1:100 in 35-mL tubes of fresh medium. Phage infection was performed by addition of 10^7^ phage after 30 min shaking. Phage-free controls were set up by adding the same volume of fresh medium instead of phage lysate. After 24 hours, supernatants and aggregates were separated as described above. To measure the resistant population of a culture, unless otherwise noted in the context, a high concentration of phage lysate was densely spread on an LM agar plate, followed by addition of the bacteria. Resistant bacteria were quantified as colony forming units on plates with phage lysate, alongside quantification of total bacteria as colony forming units on LM plates without phage lysate.

### 10-day transfer experiment in tubes and flasks with swapped growth parameters

During this experiment, we exchanged the inoculation parameters (including volume of LM medium, multiplicity of infection (MOI), temperature and shaking speeds) applied between the tube growth (5 mL, MOI 0.01, 28°C, and 160 RPM) and the flask growth (10 mL, MOI 0.1, 30°C, and 180 RPM), completely. Overnight cultures of PF4 were diluted 1:100 into 35-mL tubes containing 10 mL of fresh medium and inoculated with 1×10^7^ phages (KVP40) at an MOI of 0.1, then incubated at 30°C with shaking speed 180 RPM. In parallel, an aliquot of the same overnight PF4 cultures was diluted 1:100 into 100-mL flasks containing 5 mL of fresh medium and inoculated with 1×10^6^ KVP40 at an MOI of 0.01. These flasks were inoculated at 28°C with agitation at 160 RPM. Two sets of tube and flask cultures where the inoculation parameters were consistent with the 30-day experiments served as controls.

Every 24 hours of incubation, 1% of the cultures was transferred into fresh medium under the same condition. At designated timepoints (every three days), aliquots were taken to assess phage densities, as described above. Briefly, one mL culture samples were treated with chloroform (1%, v/v) and centrifuged (15,000g, 2 min) to isolate phages from bacteria. The lysates were diluted in phage buffer, and the phage titer was estimated by spotting 5 µL aliquots on double-layer agar plates containing *Vibrio anguillarum* PF4 as the host. The experiment continued until phages were no longer detectable.

### Coevolution experiments

Coevolution experiments in monocultures were carried out in tubes and in flasks, while experiments with the 8-strain consortium were only carried out in tubes. In the flask system for homogeneous growth, bacterial culture was transferred 1:100 into 100 mL flasks containing 10 mL fresh LM (control, three replicates) and inoculated with 1×10^7^ phage KVP40 by MOI 0.1 (coevolution, three replicates), then incubated at 30℃ with shaking speed 180rpm. Another group without phage was set up as control. In the tube cultures for heterogeneous growth, bacterial culture was transferred 1:100 into 35 mL glass tubes containing 5 mL LM (control, eight replicates) and inoculated with 1×10^6^ phage KVP40 to give MOI 0.01 (coevolution, eight replicates), then inoculated at 28℃ with shaking speed 160 RPM. For culturing of the bacterial multispecies consortium, after normalizing based on OD_600_ value of the individual monocultures, each culture of the 8 strains was mixed in equal ratios and then transferred 1:100 into 5 mL of fresh LM using 35 mL glass tubes (control, eight replicates), infected with 10^6^ phages (coevolution, eight replicates), then incubated at 30℃ with agitation 160 rpm.

Every day (after 24 hours of incubation), 1% of the cultures were transferred into fresh medium with the same condition. At the sample time points (interval two or three days), aliquots were removed to estimate bacterial and phage densities, as well as for preservation of frozen stocks for later analyses. For bacteria, aliquots were diluted in PBS and plated on LM agar (1.5%, wt/wt). For phage enumeration, 1 mL culture aliquots were treated with chloroform (1%, v/v) and centrifuged (15,000 g, 2 min) to extract phages and separate them from the bacteria. Then, lysates were diluted in phage buffer (distilled water with 5 mM Ca^2+^ and 10 mM Mg^2+^, sterilized using 0.22um filter), and the phage titer was estimated by spotting 5 µL aliquots on double-layer agar plates containing bacterial lawn of the wild-type PF4. For each time point, aliquots were preserved by freezing at −70℃ in 15% v/v glycerol.

### Phage evolution experiment with daily supply of bacterial culture

The evolution experiment was conducted under the same growth condition as used in the flasks above. The key difference was that cell-free supernatant containing phage from the previous culture was transferred into fresh medium by 1% and supplemented with fresh bacterial culture (OD_600_ ∼ 0.3). Cell-free supernatant was collected, analyzed and stored as previously described in the Coevolution section.

### Bacterial and phage isolation

To isolate bacteria, scrapes of preserved frozen samples were suspended in LM medium and subsequently spread on LM agar plates, which were then incubated overnight at 30℃. Next, colonies of different sizes were picked up and purified twice by serial streaking on LM agar plates to obtain clonal strains free of phage. Strain isolates were grown overnight at 30℃ in LM and preserved by freezing. The wild type *V. anguillarum* PF4 was used as host to isolate phages from the frozen samples using direct plating in soft agar overlays ^25^. Plaques that formed in the top agar were amplified using WT *V. anguillarum* PF4 as host. Lysates from each plaque were generated and used for the all-by-all host range cross infection matrix ^23^ as well as phage DNA extraction and sequencing in some cases.

### Estimating the proportion of resistant mutants

The susceptibility to wild type phage was tested using a cross-streak assay, which is based on assessing if bacterial growth is inhibited after streaking it left to right across a stripe of dried phage lysate on an LM agar plate ^78^. Strain isolates fell into two categories: sensitivity, indicating detectable growth inhibition on the right side of the phage lysate, compared to the left-side control; and resistance, signifying no detectable growth inhibition on the right compared to the left-side control. To screen for false negatives, killing assays were conducted in liquid medium using 96-well plates and a microtiter plate reader. In doing so, single colonies of each isolate were picked and pre-inoculated into LM for 30 min shaking at 30℃. Then, two aliquots of each culture of 20 µL from the wells were pipetted into a new 96-well plate; one was inoculated into wells containing 140 µL of LM, and the other was inoculated into wells containing 140 µL of phage lysate (1×10^8^ pfu/mL). The development of the OD_600_ values were recorded over 20 hours incubation, shaking at 30℃. For each timepoint, over 100 colonies were picked for resistant/sensitive screening and finally, 12 resistant isolates from each timepoint were randomly picked and preserved for further analyses.

### Cross-infection matrix

Resistance and infectivity phenotypes were determined by pair-wise infection ^7^. Briefly, all selected isolates were challenged with all the phage isolates, purified as described above. As a preliminary step, phage titer of each lysate as well as the ancestral KVP40 was measured by quantitative plaque assay using WT *V. anguillarum* PF4 as host. Those with a phage titer lower than 1×10^4^ PFU/mL were re-amplified to reach a higher titer. Next, all selected resistant isolates were grown overnight and 200 µL of overnight cultures of each bacterial isolate was mixed with 10 mL soft agar (0.7% v/v), and overlaid on square agar plates. Then, 5 µL drops of each lysate were spotted onto a fresh bacterial lawn and incubated at 30℃ overnight. A clearing zone was scored as a positive infection.

### Phage adsorption assay for resistant variants

To determine if selected phages were attaching to resistant isolates, we compared the number of free phage particles remaining in solution after exposure to either the resistant isolates B3, B4, B10, the WT host PF4 (positive control), and a no-host negative control. Three different colonies of each bacterial strain were each inoculated in 5 mL of LM and grown shaking at 30℃ overnight. Bacterial culture was then diluted to 1% in LM and grown shaking at 30℃ for another 3.5 hours to reach mid-exponential phase. Ten µL of Coevp10 phage lysate (1×10^9^ PFU/mL) was added to each tube and staggered in time to achieve an incubation time of 30 min for all tubes. 100 µL of the phage and bacteria mixture was then centrifuged (12,000g, 2 min) and the supernatant was supplemented with 1% chloroform to remove bacteria and any infecting or adsorbed phages. Next, 0.1 mL of a ten-fold dilution series from each replicate were quantified by double agar overlay as above. Phage adsorption was estimated by comparing the number of plaque forming units (PFUs) remaining in the cell-free supernatant to the PFUs of control tubes without bacteria ^7^.

### Passage experiment in the absence of phage

To investigate the reversal of phage resistance in the absence of phage, phage-resistant bacteria were passaged in LM media for 5 days. The wild type host PF4 was passaged alongside the resistant variants to serve as a control. Each day, 50 µL from each culture was transferred to a new tube with 5 mL of fresh media. At every transfer, a subset of each population was preserved as frozen stocks at −70℃. The experiment lasted for 5 days. At each timepoint a total of 30 colonies from each subpopulation were randomly picked for resistance/sensitivity test against the ancestral phage KVP40.

### Growth Competition

To evaluate the competitive ability of wild type PF4 and mutated variants, we conducted competition assays ^54^ between coevolved, resistant bacterial isolates (query strains: B3, B4, and B10) and the wild type host. Exponential phase cultures (OD_600_ ∼0.4-0.5) of each strain were adjusted to an OD_600_ of 0.04, and before inoculation into fresh medium either in single culture or mixed in a 1:1 ratio with the competitor, aliquots of each single culture were diluted and plated to enumerate the initial bacterial densities. Then, three rounds of transfers into fresh medium were carried out, in which samples were collected with OD_600_ between 0.6 (first time) to 0.9 (last two times) and were diluted and plated to obtain final densities. For CFU quantification, the collected samples were immediately diluted with PBS (10ul sample + 990 µl PBS) and mixed thoroughly, subsequently passed through a serial 10-fold dilution series. To quantify the phage-resistant proportion of the cultures, unless otherwise noted in the context, colonies formed on LM agar plates ^7^, which contained a high concentration of KVP40 phage lysate on their surface were considered resistant and compared to the number of colonies formed on LM agar plates without phage.

### Biofilm assay

Biofilm assays were conducted in 96-well microplates following a method described previously by Faizan *et al*. ^79^, with some modifications. Briefly, overnight cultures of each resistant isolate were inoculated in 96-well plates by 1% with 150 µL fresh LM as the inoculum volume. The microtiter plates were incubated under static conditions at 30℃ for 24 hours for biofilm formation. The supernatant was removed from all wells using a multichannel pipette and the plates were rinsed with 150 µl of fresh LM three times. Biofilms were stained with 0.5% (w/v) crystal violet (CV), 160 µl per well, and incubated for 30 minutes at room temperature. The excess CV was rinsed off with PBS. After solubilizing CV using 200 µl of 96% ethanol, plates were incubated with agitation for 30 min and absorbance was measured at 595 nm. When absorbance values exceeded 2, CV was diluted with ethanol. In each plate, wells inoculated with wild type PF4 and fresh LM medium served as the reference strain and the negative control, respectively.

### Liquid culture phage infection

Overnight cultures were diluted 1:100 in LM supplemented with 10mM MgSO_4_, and grown at 30℃ with shaking for 30 min. A calculated volume of phage stock (5.0 ×10^7^ PFU) was added to each culture and 150 µL was seeded into each well of a 96-well plate. OD_600_ was measured every 5 min in a microplate reader at 30℃ with shaking.

### DNA sequencing

Single colonies were inoculated in 35-mL glass tubes containing 5 mL LM and grown shaking at 30℃ overnight. The fresh culture was pelleted by centrifugation at 5,000 g for 10 min and then immediately processed for DNA extraction. The Qiagen Genomic Tip 500x kit was used following the manufacturers guidelines, and the final DNA was collected by spooling using a glass rod rather than centrifugation to avoid shearing. DNA samples were stored in 70 µL of elution buffer at 4℃ for 50 hours to allow for full resuspension before sequencing at BGI (www.bgi.com). Phage genomic DNA was extracted using Monarch HMW DNA extraction kit (NEB #T3060) following the manufacturers guidelines. Bacterial and phage DNA concentration purity were measured with NanoDrop and the final concentration was determined by QUBIT. Sequencing library construction and sequencing was carried out by BGI (DNBseq platform) to produce 150-bp paired-end reads. In parallel, to obtain closed genomes for comparative genomic analysis, the chromosome DNA of the wild type *V. anguillarum* PF4 was also sequenced on Oxford Nanopore sequencer for long reads. DNA sample was prepared for nanopore sequencing according to recommendations by ONT with a rapid barcoding sequencing kit (SQK-RBK004; ONT) and sequenced on an ONT GridION system.

### Sequencing analysis

Illumina reads were filtered to remove reads contaminated by the Nextera adapter or low-quality bases (>2 bases with a Phred Score of <20) ^80^. Trimmomatic was used to clean and trim the sequencing reads yielding an average sequencing read length of 150 bp with depth around 100, before mapping them on the reference genome using Bowtie2(v2.5) ^81^.

For bacterial reference genome preparation, a hybrid *de novo* assembly pipeline with Illumina short reads and nanopore long reads was designed for data processing. In brief, long reads were filtered and trimmed with NanoPack(v1.10.0) ^82^ after basecalling with Guppy(v3) ^83^, and then assembled using Minimap2(v2.28) ^84^ and Nanopolish(v0.14.0) ^85^ to create a set of reference genomes. The quality and completeness of the reference genomes were evaluated using QUAST(v5.2) ^86^, and then the best genome was selected and used as a reference in the final assembly. In the final assembly stage, both sets of processed reads, together with the best reference assembly, were used as inputs for the Unicycler ^87^ assembler.

Genomes of the wild type *V. anguillarum* PF4 and phage KVP40, were annotated using Prokka (v1.14.5)^88^. SNP analysis was performed using an in-house analysis pipeline. Briefly, reads were aligned to a reference genome (PF4 for bacteria and KVP40 for phages) using Bowtie2(v2.5) ^81^ with a maximum of 3 mismatches per read. Base calling was done using SAMtools(v1.20.0) ^89^ and BCFtools(1.20.0) ^90^, and a genome position was determined as a SNP when more than a single allele was identified among all isolates with a quality threshold of FQ< −80. In parallel, set to polymorphism mode, the computational analysis pipeline Breseq(v0.38.3) ^91^ was also adopted for insertions and deletion identification with the same reference genomes. To identify amplification and additional deletions, the genome coverage of each isolate was normalized in two steps: first, the coverage values of each isolate were divided by the isolate median coverage, and then the normalized values were further divided by the corresponding normalized base pair coverage value of the ancestral genome. Areas with high or low coverage were identified manually. Phylogeny was based on the identified SNPs and was determined by the Harvest parsnp algorithm(v2.0.5) ^92^, which was carried out as unrooted parsimony. Adaptive evolution significance was evaluated by randomly distributing the total number of intragenic SNPs across all ORFs, while accounting for mutations that appeared in any of the isolates and preserving transition/transversion ratios.

### Data analysis

All statistical analyses were conducted in the R open-source software (R v4.3.3). All statistical tests were two-sided and differences were considered statistically significant if the *P* value <0.05. In particular, the stats packages were used to run all ANOVAs and Mann-Whitney U test (using the lm and wilcox test functions).

## Data availability

Genome sequencing data is deposited in the public SRA database, with accession code PRJNA1090900: SAMN40578671 - SAMN40578683 for phage sequencing data and SAMN40577831 - SAMN40577841 for bacterial genomic sequencing data. All other datasets generated and analyzed during the current study were deposited in the GitHub repository with https://github.com/lingc31415/phage-bacteria-interaction-in-coevolutionary-scope, together with the custom scripts.

## Code availability

The present study mentioned tools used for the data analysis were applied with default parameters unless specified otherwise.

## Acknowledgments

We thank B. K. Gummesson. for expert technical assistance; Y. F. Ma and Z. P. Huang for generously providing the strains used in the community development process; L. Krych, D. V. Stefanova, and S. J. Traving for help with nanopore sequencing. This study was financially supported by the European Union under the Horizon Europe Programme, Grant Agreement No. 101084204 (Cure4Aqua), by the Danish Innovation Fund project AQUAPHAGE, and Danish National Research Foundation through the Danish Center for Hadal Research HADAL, Grant No. DNRF145 to MM, and Novo Nordisk Foundation grant number NNF22OC0079521 to SLS.

## Author contributions

L.C., M.M., and S.L.S. conceived and designed the research. L.C performed experiments and analysis and L.C., M.M, and S.L.S. interpreted the results. L.C., S.L.S., and M.M., prepared the manuscript. All the authors reviewed the results and approved the final version of the manuscript.

