## Supplementary material for "Spatial heterogeneity influences phage-host co-existence dynamics and phage resistance development in *Vibrio anguillarum*"

Submitted to BioRXiv on December 28, 2024, by Sine Lo Svenningsen

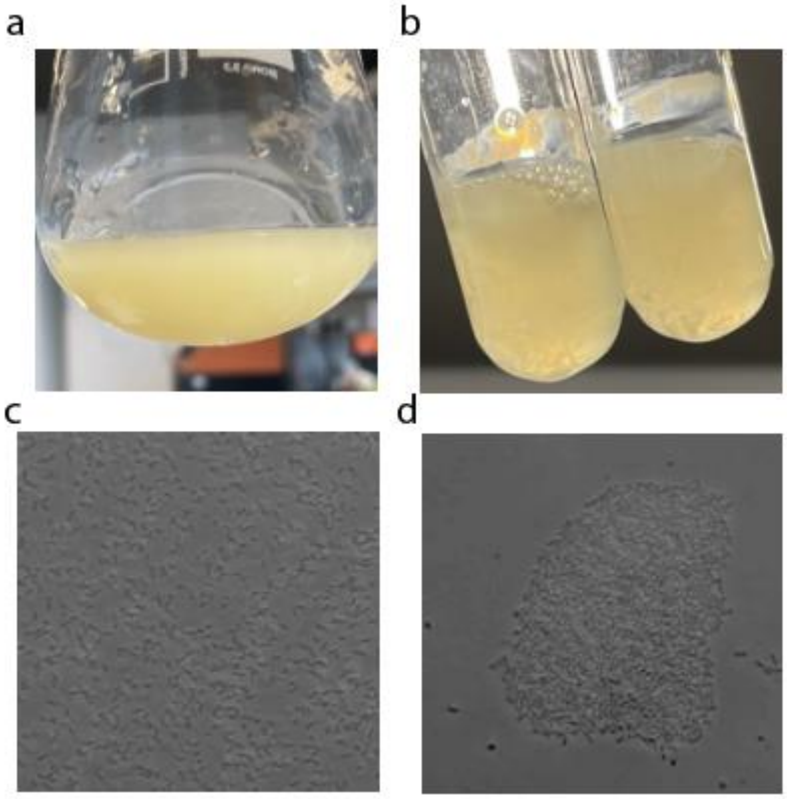


**Fig. S1. Bacterial phenotypic responses to divergent growth conditions**

Aliquots of a mixture of phage and bacteria were inoculated in both flasks (a) and tubes (b). Cultures were harvested and subjected to microscopy after incubation for 24 hours growth at 30℃. Morphological visualization of *V. anguillarum* PF4 aliquots after 24-hour culturing in flasks (**c**) and aggregates in tubes (**d**).

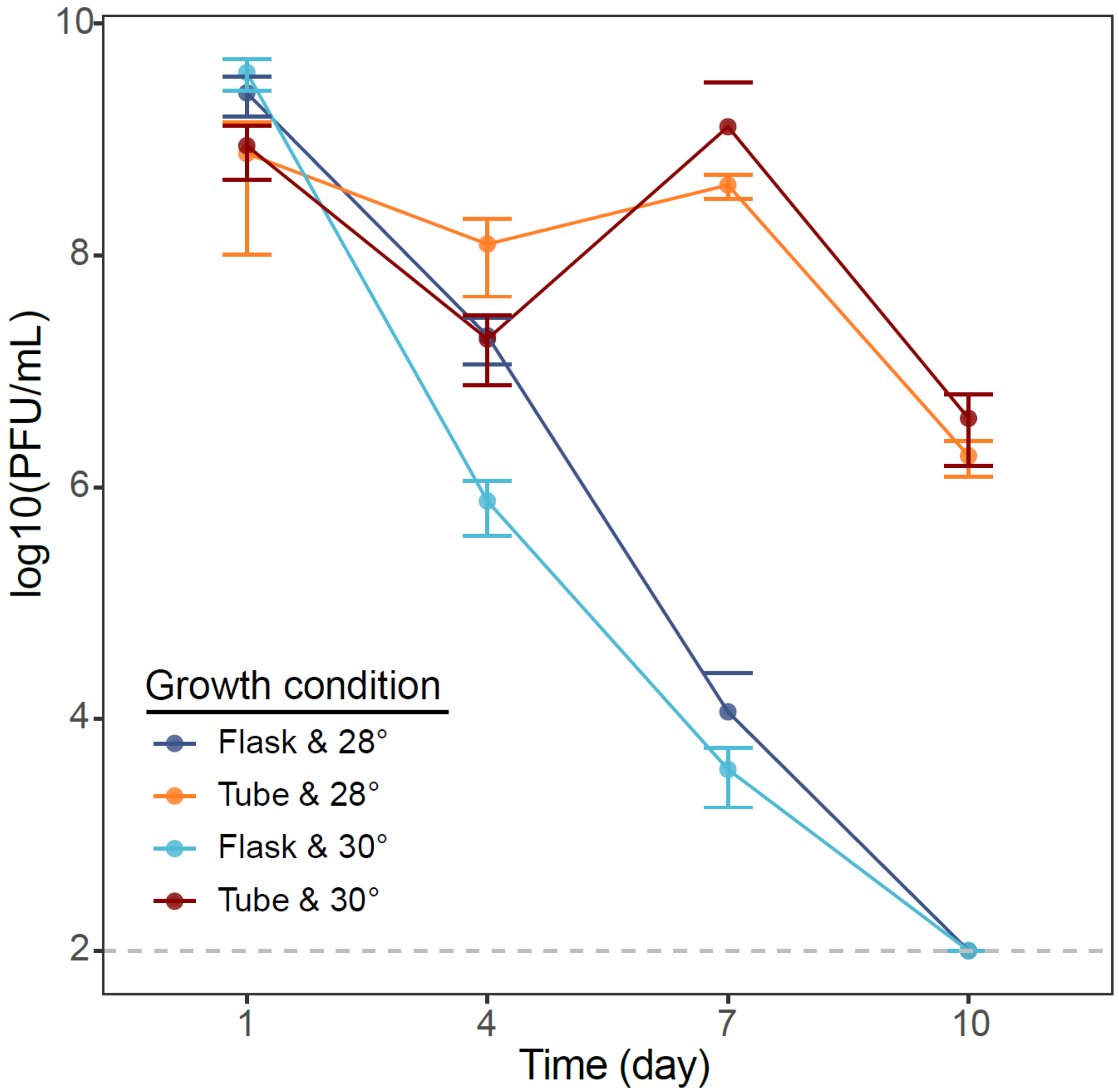


**Fig. S2. Phages persist longer in tubes than flasks independent of specific growth parameters** Phage populations from cultures incubated in tubes are represented in orange (5 mL volume, initial MOI 0.01, 28°C, and 160 RPM shaking speed) and in dark red (10 mL volume, initial MOI 0.1, 30°C, and 180 RPM shaking speed). Phage populations from cultures incubated in flasks are shown in dark blue (5 mL volume, initial MOI 0.01, 28°C, and 160 RPM shaking speed) and light blue (10 mL volume, MOI 0.1, 30°C, and 180 RPM shaking speed). Phage populations were quantified in triplicate for each condition and time point.

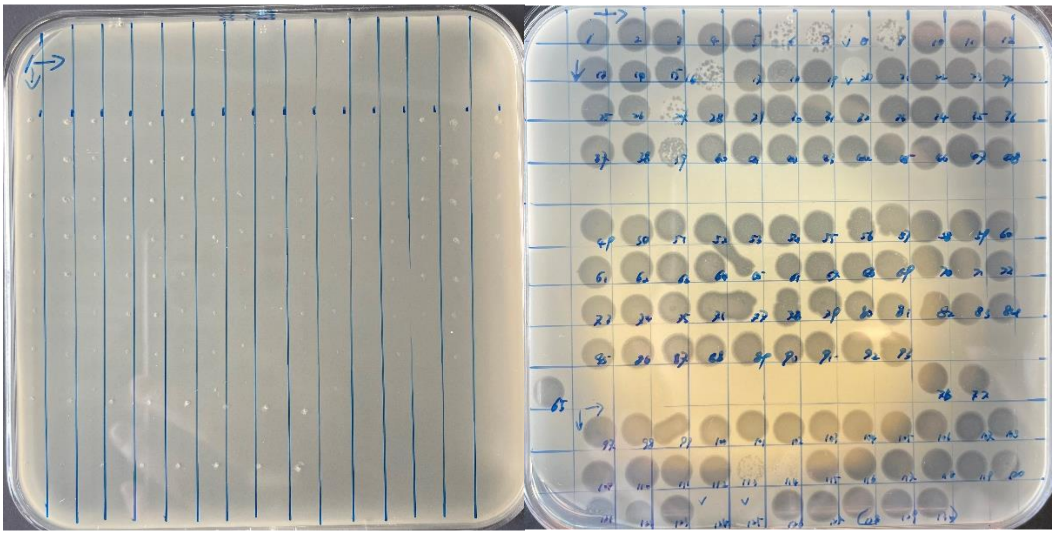

**Fig. S3. Pair-wise all-by-all cross-infection matrix**

The left side shows an example of a spot test of 196 phages against one of the bacterial isolates that are resistant to the ancestral phage KVP40. The right side shows a spot test of the phage isolates on the wild type PF4 strain for reference. Phages were spotted onto lawns of each of the bacterial strains and incubated overnight at 30 degrees. The transparent clear zones seen on the plate on the right indicate infection and host cell lysis. No infections were detected by any of the 196 phages on any of the 120 KVP40-resistant isolates tested in the experiment.

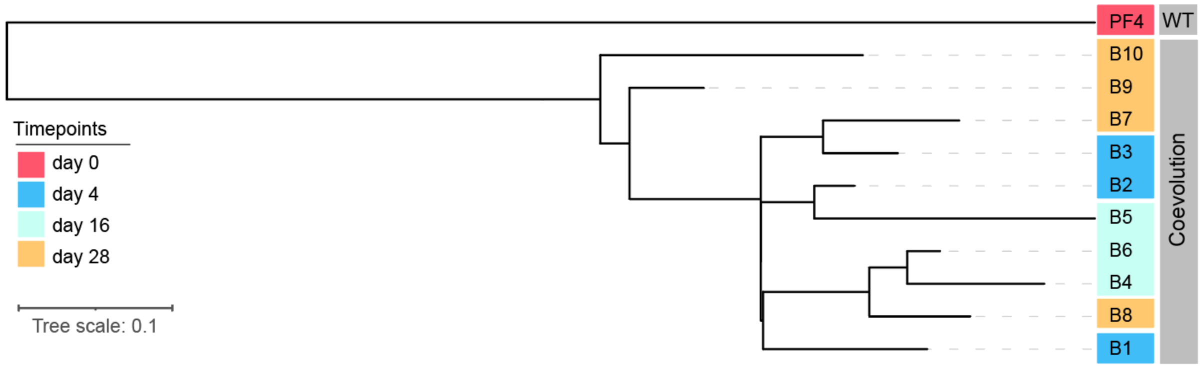


**Fig. S4. Phylogeny of sequenced bacterial isolates from the coevolution experiment in tubes**

The unrooted tree was constructed with whole genome SNPs from Parsnp [1] program and optimized by IQ-tree [2].

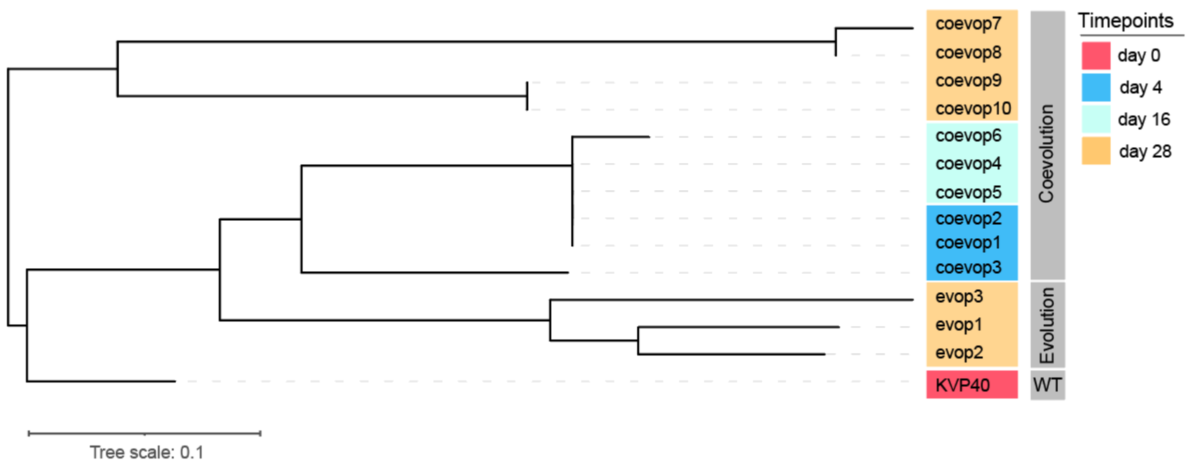


**Fig. S5. Phylogenetic relationship among the phage isolates from the coevolution experiment in tubes**

The relationship was estimated by whole genome comparison. The unrooted tree was constructed with whole genome SNPs from Parsnp [1] program and optimized by IQ-tree [2].

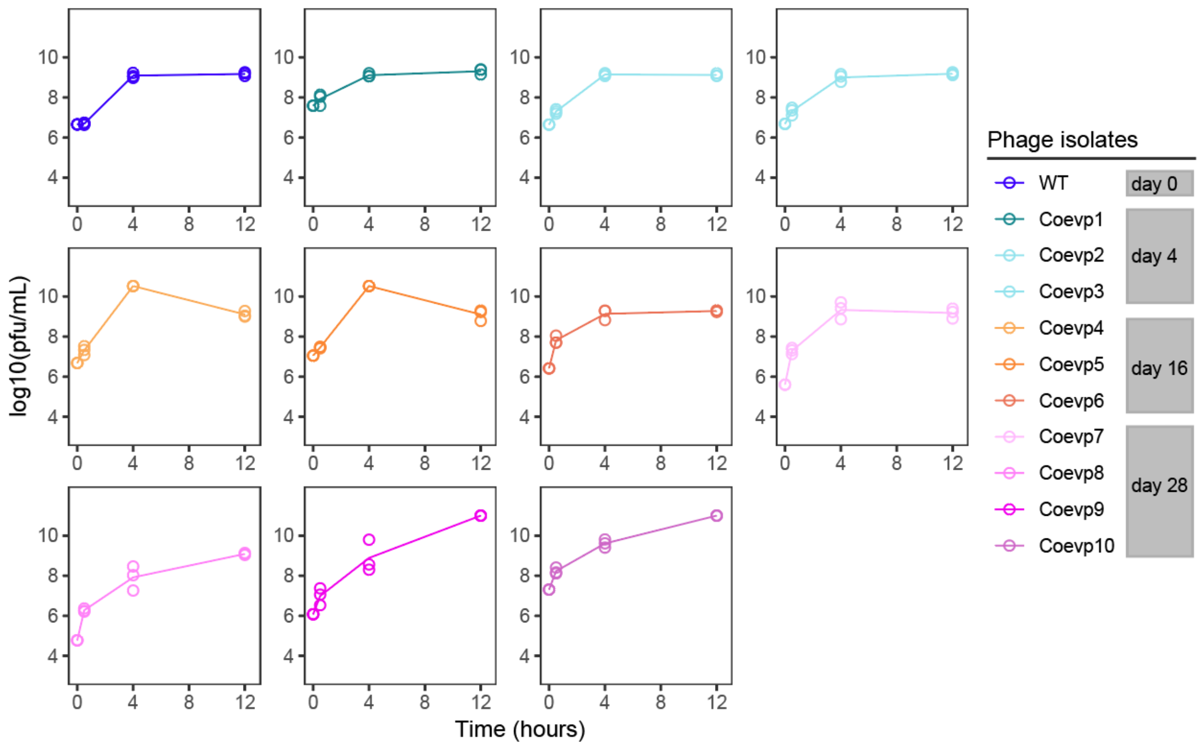


**Fig. S6. Infective traits of the genome-sequenced coevolved phages**

Phage infection assays with phage isolates from the coevolution experiments in tubes. The infections were initiated at MOI ~ 0.01. The ancestral phage was used as control (blue). Each experiment had three biological replicates, all of which are shown on the graphs.

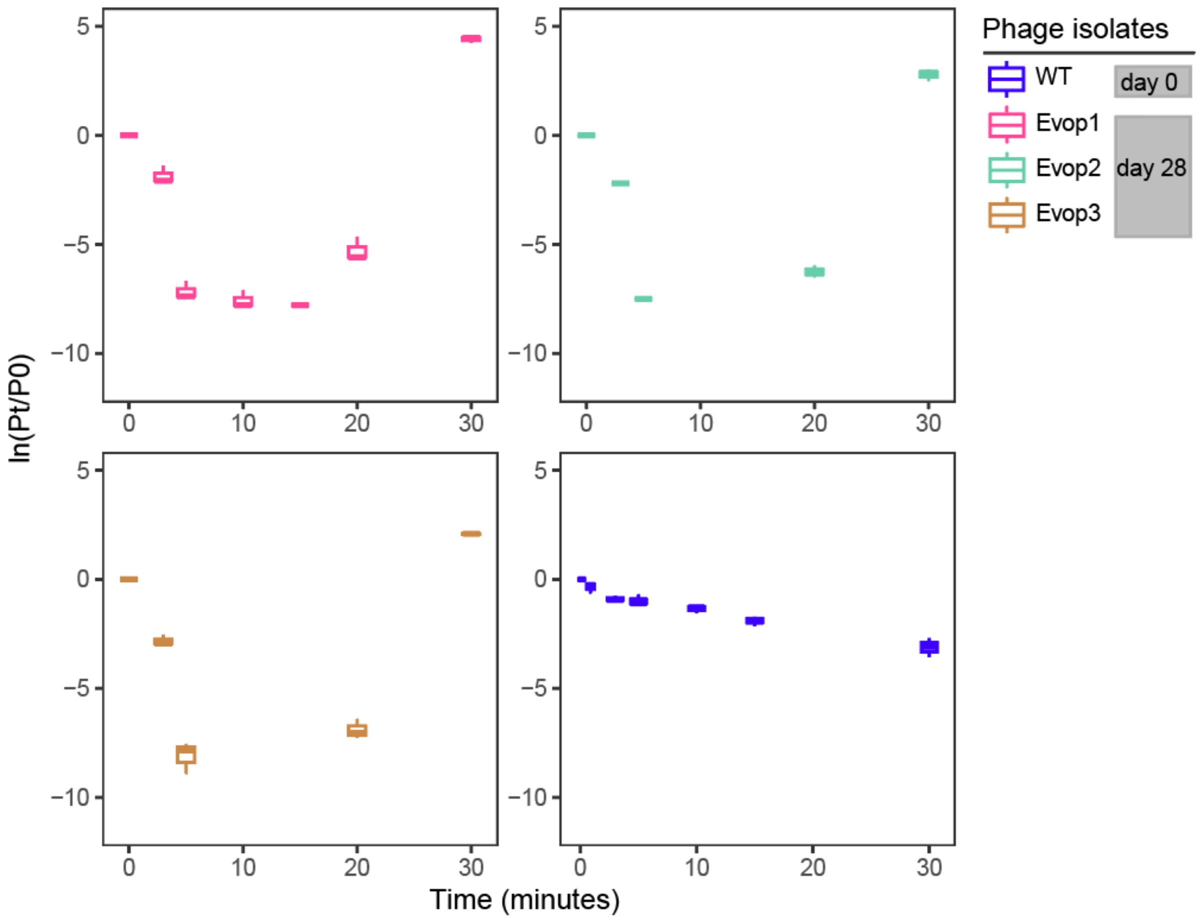


**Fig. S7. Phenotypic variation of three trained phages**

Early infection dynamics of three phages isolated from the phage evolution experiment, where phages were transferred to fresh susceptible hosts every 24 hours. At time 0, phages were added to *V. anguillarum* PF4 at MOI 0.01, and phage numbers were monitored for 30 minutes with seven timepoints. Each phage isolate was cultured in three biological replicates. Pt indicates the phage titer (PFU/mL) at the indicated time point, P0 indicates the phage titer (PFU/mL) at time 0.

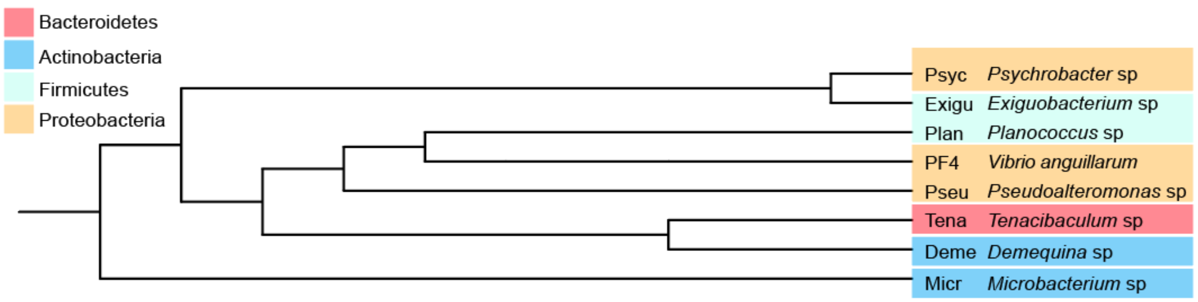


**Fig. S8. Phylogenetic tree of bacterial strains comprising the multispecies community**

Phylogenetic analysis was performed using a concatenated alignment of 16S rRNA genes. The Neighbor-joining tree was generated with 1000 bootstraps. Branches are marked in different colors according to the phylum.

**Table S1. Mutations in bacterial genomes isolated from the coevolution experiment in tubes**

Mutations were detected by combining Illumina short read sequencing and Nanopore long read sequencing as input into the computational pipeline Breseq [3]. The position of the mutation in the genome, the type of mutation, the position of the mutation relative to annotated genes, and the description of the mutated gene are displayed in the table. A filled gray square indicates the presence of the mutation in the genome of bacterial isolates B1-B10. Annotations in blue, green, and red text correspond to nonsynonymous, synonymous, and stop codon mutations, respectively.

| Seq id | Position | Mutation | day4 | | | day16 | | | day28 | | | | Annotation | Description |
|---|---|---|---|---|---|---|---|---|---|---|---|---|---|---|
| | | | B1 | B2 | B3 | B4 | B5 | B6 | B7 | B8 | B9 | B10 | | |
| contig_2 | 51,564 | (T)$_{14\to16}$ | ■ | ■ | | | | ■ | ■ | ■ | ■ | | intergenic (+2092/+81) | Epoxyqueuosine reductase/Oligoribonucleas e |
| contig_2 | 51,564 | (T)$_{14\to17}$ | | | ■ | | ■ | | | | | ■ | intergenic (+2092/+81) | Epoxyqueuosine reductase/Oligoribonucleas e |
| contig_2 | 188,914 | T→G | | | | | | | ■ | | | | D169A (GAC→GCC) | Cell division ATP-binding protein FtsE |
| contig_2 | 189,302 | C→T | | | | | | | | ■ | | | G40S (GGT→AGT) | Cell division ATP-binding protein FtsE |
| contig_2 | 298,497 | +107 bp | | | | | | | | ■ | | ■ | intergenic (+2348/-9702) | Protoporphyrinogen IX dehydrogenase [menaquinone]/Putative fluoride ion transporter CrcB |
| contig_2 | 821,138 | (A)$_{8\to9}$ | | | | | | | | | ■ | ■ | coding (11/1731 nt) | Peptidoglycan D,D-transpeptidase FtsI |
| contig_2 | 844,740 | T→A | ■ | ■ | ■ | ■ | | ■ | ■ | ■ | ■ | ■ | C580S (TGT→AGT) | Carbamoyl-phosphate synthase large chain |
| contig_2 | 937,647 | Δ1 bp | | | | | | ■ | | | ■ | ■ | coding (760/807 nt) | Nucleoside-specific channel-forming protein Tsx |
| contig_2 | 937,816 | G→C | | ■ | | ■ | | | | | | | Y197* (TAC→TAG) | Nucleoside-specific channel-forming protein Tsx |
| contig_2 | 938,012 | (GTTAAC ACCGAAA GTTGTAC TTGTTG)$_{1\to2}$ | | | | | | | ■ | | | | coding (395/807 nt) | Nucleoside-specific channel-forming protein Tsx |
| contig_2 | 938,291 | Δ90 bp | ■ | | ■ | | ■ | | | ■ | | | coding (27-116/807 nt) | Nucleoside-specific channel-forming protein Tsx |

| | | | | | | | | | | | | | | |
|---|---|---|---|---|---|---|---|---|---|---|---|---|---|---|
| contig_2 | 1,309,009 | C→G | | | | | | | | | | | T307R (ACG→AGG) | Glutathionyl-hydroquinone reductase YqjG |
| contig_2 | 2,786,453 | +107 bp | | | | | | | | | | | intergenic (+4206/–) | Na(+)/H(+) antiporter NhaP/– |
| contig_4 | 157,276 | T→G | | | | | | | | | | | I41M (ATT→ATG) | hypothetical protein |
| contig_4 | 227,660 | T→C | | | | | | | | | | | A59A (GCA→GCG) | hypothetical protein |
| contig_4 | 238,726 | Δ1,093 bp | | | | | | | | | | | | unknown |

**Table S2 (part 1). Mutations in phage isolated from both the coevolution experiments in tubes and the evolution experiments in flasks.** Mutations were detected by combining Illumina short read sequencing and Nanopore long read sequencing as input into the computational pipeline Breseq [3]. A filled gray square indicates the presence of the mutation in the genome of phage isolates. Annotations in blue, green, and red text correspond to nonsynonymous, synonymous, and nonsense mutations, respectively.

| Position | Mutation | Coevolution (Coevo-) | | | | | | | | | | Evolution | | | Annotation | Description |
|---|---|---|---|---|---|---|---|---|---|---|---|---|---|---|---|---|
| | | day4 | | | day16 | | | day28 | | | | day28 | | | | |
| | | p1 | p2 | p3 | p4 | p5 | p6 | p7 | p8 | p9 | p10 | Evop1 | Evop2 | Evop3 | | |
| 32,988 | T→G | ■ | ■ | ■ | ■ | ■ | ■ | ■ | ■ | ■ | ■ | ■ | ■ | ■ | E426A (GAG→GCG) | hypothetical protein |
| 76,168 | C→T | | | | | | | | | | | | ■ | ■ | A12V (GCA→GTA) | hypothetical protein |
| 76,355 | C→A | | | | | | | ■ | ■ | | | | | | S74R (AGC→AGA) | hypothetical protein |
| 80,361 | C→A | ■ | ■ | ■ | ■ | ■ | ■ | | | | | | | | A153E (GCA→GAA) | DNA methyltransferase |
| 88,016 | G→A | | | | | | | ■ | ■ | | | | | | intergenic (-24/-229) | hypothetical protein/hypothetical protein |
| 93,719 | A→T | | | | | | | | | | | ■ | | | D93V (GAT→GTT) | hypothetical protein |
| 93,729 | A→C | | | | | | | | | ■ | ■ | | | | K96N (AAA→AAC) | hypothetical protein |
| 97,393 | Δ9 bp | | | | | | | | | | | ■ | ■ | ■ | intergenic (+107/-52) | hypothetical protein/hypothetical protein |
| 98,230 | Δ8 bp | | | | | | | | | | | ■ | ■ | ■ | coding (24-31/336 nt) | hypothetical protein |
| 98,314 | 86 bp→65 bp | | | | | | | | | | | | ■ | ■ | coding (108-193/336 nt) | hypothetical protein |
| 98,390 | Δ10 bp | | | | | | | | | | | ■ | | | coding (184-193/336 nt) | hypothetical protein |
| 98,390 | 38 bp→19 bp | | | | | | | ■ | ■ | | | | | | coding (184-221/336 nt) | hypothetical protein |
| 98,483 | Δ10 bp | | | | | | | | | | | ■ | ■ | ■ | coding (277-286/336 nt) | hypothetical protein |
| 98,508 | Δ10 bp | | | | | | | ■ | ■ | | | | | | coding (302-311/336 nt) | hypothetical protein |
| 98,808 | C→A | | | | | | | ■ | ■ | | | | | | intergenic (+95/-380) | hypothetical protein/hypothetical protein |
| 98,809 | C→T | ■ | ■ | ■ | ■ | ■ | ■ | | | ■ | ■ | | | | intergenic (+96/-379) | hypothetical protein/hypothetical protein |
| 98,809 | 1 bp→99 bp | | | | | | | | | | | | | ■ | intergenic (+96/-379) | hypothetical protein/hypothetical protein |
| 98,812 | G→T | | | | | | | ■ | ■ | | | | ■ | | intergenic (+99/-376) | hypothetical protein/hypothetical protein |
| 98,854 | Δ8 bp | | | | | | | | | | | ■ | | | intergenic (+141/-327) | hypothetical protein/hypothetical protein |
| 98,862 | +76 bp | | | | | | | | | | | | ■ | | intergenic (+149/-326) | hypothetical protein/hypothetical protein |
| 98,870 | Δ10 bp | | | | | | | ■ | ■ | | | | | | intergenic (+157/-309) | hypothetical protein/hypothetical protein |
| 98,893 | Δ13 bp | | | | | | | | | | | | | ■ | intergenic (+180/-283) | hypothetical protein/hypothetical protein |
| 98,894 | A→G | ■ | ■ | ■ | ■ | ■ | ■ | | | | | | | | intergenic (+181/-294) | hypothetical protein/hypothetical protein |
| 98,896 | Δ1 bp | | | | | | | | | ■ | ■ | | | | intergenic (+183/-292) | hypothetical protein/hypothetical protein |
| 98,897 | Δ1 bp | | | | | | | | | | | ■ | | | intergenic (+184/-291) | hypothetical protein/hypothetical protein |
| 98,905 | 3 bp→AAA | | | | | | | | | | | ■ | | | intergenic (+192/-281) | hypothetical protein/hypothetical protein |
| 98,906 | 2 bp→AA | | | | | | | | | | | | ■ | | intergenic (+193/-281) | hypothetical protein/hypothetical protein |
| 98,944 | +CACCTG TGTAAT | | | | | | | ■ | | | | | ■ | | intergenic (+231/-244) | hypothetical protein/hypothetical protein |
| 98,951 | A→T | | | | | | | | | | | | ■ | | intergenic (+238/-237) | hypothetical protein/hypothetical protein |
| 98,961 | Δ14 bp | | | | | | | | | | | | ■ | | intergenic (+248/-214) | hypothetical protein/hypothetical protein |

| | | | | | | | | | | | | | | | | |
|---|---|---|---|---|---|---|---|---|---|---|---|---|---|---|---|---|
| 98,984 | (42 bp)$_{1\rightarrow2}$ | | | | | | | | | | | | | | intergenic (+271/-204) | hypothetical protein/hypothetical protein |
| 101,825 | G→A | | | | | | | | | | | | | | C32Y (TGT→TAT) | hypothetical protein |

**Table S2 (part 2). Mutations in phage isolated from both the coevolution experiments in tubes and the evolution experiments in flasks.**

| Position | Mutation | Coevolution (Coevo-) | | | | | | | | | | Evolution | | | Annotation | Description |
|---|---|---|---|---|---|---|---|---|---|---|---|---|---|---|---|---|
| | | day4 | | | day16 | | | day28 | | | | day28 | | | | |
| | | p1 | p2 | p3 | p4 | p5 | p6 | p7 | p8 | p9 | p10 | Evop1 | Evop2 | Evop3 | | |
| 102,038 | G→C | | | | | | | | | | | | | | G103A (GGT→GCT) | hypothetical protein |
| 105,825 | A→G | | | | | | | | | | | | | | Y177C (TAC→TGC) | thymidine kinase |
| 119,071 | C→A | | | | | | | | | | | | | | L53I (CTT→ATT) | hypothetical protein |
| 120,044 | Δ5,711 bp | | | | | | | | | | | | | | | *[KVP40.0227], KVP40.0228, KVP40.0229, KVP40.0230, KVP40.0231, KVP40.0232, KVP40.0233, KVP40.0234, KVP40.0235, KVP40.0236, KVP40.0237, KVP40.0238, KVP40.0239, [KVP40.0240]* |
| 124,366 | +GA | | | | | | | | | | | Δ | Δ | Δ | coding (314/633 nt) | DNA endonuclease V |
| 124,531 | T→C | | | | | | | | | | | Δ | Δ | Δ | L160P (CTG→CCG) | DNA endonuclease V |
| 125,065 | Δ11 bp | | | | | | | | | | | Δ | Δ | Δ | coding (384-394/615 nt) | hypothetical protein |
| 131,622 | C→G | | | | | | | | | | | | | | S133R (AGC→AGG) | hypothetical protein |
| 146,183 | T→A | | | | | | | | | | | | | | V52D (GTT→GAT) | hypothetical protein |
| 161,840 | Δ360 bp | | | | | | | | | | | | | | intergenic (-852/+2169) | hypothetical protein/long tail fiber protein distal subunit |
| 162,742 | C→T | | | | | | | | | | | | | | intergenic (-1754/+1626) | hypothetical protein/long tail fiber protein distal subunit |
| 163,008 | T→C | | | | | | | | | | | | | | intergenic (-2020/+1360) | hypothetical protein/long tail fiber protein distal subunit |
| 163,136 | T→C | | | | | | | | | | | | | | intergenic (-2148/+1232) | hypothetical protein/long tail fiber protein distal subunit |
| 181,479 | (GGATGCTTACG)$_{1\rightarrow2}$ | | | | | | | | | | | | | | intergenic (-266/+51) | tRNA-OTHER/hypothetical protein |
| 195,183 | Δ11 bp | | | | | | | | | | | | | | coding (635-645/876 nt) | hypothetical protein |
| 195,689 | G→C | | | | | | | | | | | | | | P47A (CCA→GCA) | hypothetical protein |
| 198,605 | T→A | | | | | | | | | | | | | | V44D (GTC→GAC) | baseplate wedge subunit |
| 199,210 | T→C | | | | | | | | | | | | | | Y246H (TAT→CAT) | baseplate wedge subunit |
| 199,308 | T→A | | | | | | | | | | | | | | D278E (GAT→GAA) | baseplate wedge subunit |
| 201,293 | G→T | | | | | | | | | | | | | | S940I (AGT→ATT) | baseplate wedge subunit |
| 202,109 | G→A | | | | | | | | | | | | | | D49N (GAT→AAT) | baseplate wedge subunit |
| 202,385 | G→T | | | | | | | | | | | | | | D141Y (GAC→TAC) | baseplate wedge subunit |
| 203,368 | G→C | | | | | | | | | | | | | | G115A (GGC→GCC) | baseplate wedge tail fiber protein connector |
| 205,929 | G→A | | | | | | | | | | | | | | E638K (GAA→AAA) | baseplate wedge subunit |
| 206,065 | A→G | | | | | | | | | | | | | | E683G (GAG→GGG) | baseplate wedge subunit |
| 206,848 | G→T | | | | | | | | | | | | | | S201S (TCG→TCT) | baseplate wedge subunit |
| 210,797 | A→G | | | | | | | | | | | | | | E198G (GAA→GGA) | fibritin neck whisker |
| 211,070 | T→A | | | | | | | | | | | | | | V289D (GTT→GAT) | fibritin neck whisker |
| 211,742 | C→A | | | | | | | | | | | | | | T513K (ACA→AAA) | fibritin neck whisker |
| 218,563 | T→G | | | | | | | | | | | | | | S158A (TCT→GCT) | tail sheath |
| 221,313 | C→T | | | | | | | | | | | | | | P202S (CCG→TCG) | portal protein |
| 221,828 | T→A | | | | | | | | | | | | | | D373E (GAT→GAA) | portal protein |
| 227,592 | C→A | | | | | | | | | | | | | | A110E (GCG→GAG) | hypothetical protein |
| 227,692 | G→A | | | | | | | | | | | | | | Q22Q (CAG→CAA) | hypothetical protein |
| 230,101 | T→C | | | | | | | | | | | | | | T365A (ACT→GCT) | hypothetical protein |

| Position | Mutation | Coevolution (Coevo-) | | | | | | | | | | Evolution | | | Annotation | Description |
| --- | --- | --- | --- | --- | --- | --- | --- | --- | --- | --- | --- | --- | --- | --- | --- | --- |
| | | day4 | | | day16 | | | day28 | | | | day28 | | | | |
| | | p1 | p2 | p3 | p4 | p5 | p6 | p7 | p8 | p9 | p10 | Evop1 | Evop2 | Evop3 | | |
| 235,642 | G→A | | | | | | | | | | | | | | G108D (GGT→GAT) | DNA helicase |
| 240,211 | C→T | | | | | | | | | | | | | | G269D (GGT→GAT) | hinge connector of long tail fiber protein distal connector |
